# Elevated precontractile calcium set point is sustained by limited PMCA1 export in dystrophic muscle

**DOI:** 10.64898/2026.11.05.764057

**Authors:** D Marchiafava, K Hughes-Wiles, W Stein, X Dai, P Jazireian, A Stanley-Copeland, J Ohana, MF Engelke, AG Vidal-Gadea

## Abstract

Duchenne muscular dystrophy is caused by loss of dystrophin and marked by elevated muscle calcium, usually attributed to contraction-induced membrane damage. Whether calcium rises before muscle can contract, and what sustains it, is unresolved. In *Caenorhabditis elegans* embryos and patient-derived human myoblasts, dystrophic muscle carries an elevated resting calcium set point before contraction is possible. Embryonic calcium is elevated 400 minutes after fertilization, before movement is detectable or the neuromuscular junction forms. Silencing the plasma membrane calcium pump gene *mca-1* raises calcium in wild-type but not dystrophic animals, whereas silencing its paralogue raises it in both: the limitation is specific to that export route. Dystrophic human myoblasts have less pump; pump inhibition or silencing increases calcium preferentially in healthy cells and raising pump expression lowers calcium most in dystrophic cells. Constrained calcium export is an early, continuing determinant of the dystrophic set point.

## Introduction

Duchenne muscular dystrophy (DMD) is an X-linked recessive disorder affecting approximately 1 in 5,000 males(*1*). DMD arises from loss of function mutations in the *DMD* gene encoding dystrophin, which connects the cells’ actin cytoskeleton to the extracellular matrix through the dystrophin glycoprotein complex (DGC)(*1*, *2*). Mutations of the dystrophin gene result in DGC disruption, cell membrane instability, and pathological changes including increased sarcoplasmic calcium(*3–6*). These phenotypes lead to continued muscle weakness and, ultimately, contraction failure with lethal outcome.

Calcium is a critical intracellular messenger involved in contraction, metabolism, and signal transduction(*7*). Calcium levels are controlled by a diverse array of channels and pumps that regulate spatial and temporal calcium signals with high precision(*8*, *9*). In dystrophin-deficient muscle, this regulatory balance is lost early in childhood, leading to elevated calcium levels, hypercontractility, activation of proteases, mitochondrial dysfunction, and ultimately muscle necrosis(*1*). However, the molecular origins of early calcium dysregulation in dystrophic muscle remain unclear. Does early calcium elevation arise from defects in calcium handling and thus independently of mechanical stress during muscle contraction, or is it a consequence of active force generation in muscles that tear cell membranes?

These are distinctions that carry substantial conceptual weight. If calcium becomes elevated before contractile loading, i.e., not as a mere consequence of membrane tearing, Duchenne muscular dystrophy might be as much a developmental disorder as it is a progressive condition. This could impact treatment options and recommendations for occupational and physical therapy. It also holds conceptual significance as the time point at which calcium dysregulation occurs may provide clues about the molecular origins of a major contributor to the phenotype of this disease.

In mature muscle, calcium homeostasis reflects the coordinated action of many systems, including plasma membrane channels, pumps, and intracellular stores such as the sarcoplasmic reticulum (SR)(*10*). The relative importance of these systems changes as myogenic cells differentiate and muscle contraction becomes more prevalent and components such as the sarco/endoplasmic reticulum Ca²⁺-ATPase (SERCA) and ryanodine receptor mediated flux take on a larger share of calcium control(*11*, *12*). Accordingly, many studies have pointed towards contributions of contraction-mediated calcium dysregulation to the sustained shift in basal cytosolic calcium that is observed in DMD. Nevertheless, the initiating causes of the calcium imbalance remain elusive(*13*). If calcium imbalance were to stem from pre-contractile developmental stages, the mechanistic underpinnings driving it would be very distinct. In pre-contractile muscle cells the muscle-specific architecture and its contraction machinery have not reached full functionality yet(*14*, *15*). As in most non-contractile eukaryotic cells, calcium extrusion through the cell membrane is a core requirement for maintaining low resting cytosolic calcium at this developmental stage(*16*).

At early stages of myogenesis, the full DGC is not yet assembled. Cellular processes that depend on interactions between dystrophin and the sarcolemma and other components of the DGC are not yet functional. The plasma membrane calcium ATPases (PMCAs) PMCA1 and PMCA4 are broadly expressed in tissues, and most abundantly expressed in skeletal muscle. These proteins function to maintain low resting cytoplasmic calcium levels independently of the contractile apparatus that assembles after dystrophin expression.

Because PMCA1 and 4-mediated calcium extrusion is present before cells commit to differentiation, and as a result contraction, PMCAs may be plausible candidates for contributors to early calcium dysregulation in dystrophic muscle during myogenesis(*17–19*).

Mammalian DMD models such as mice and canines capture many features of DMD but present practical challenges for longitudinal *in vivo* imaging of calcium dynamics during early myogenesis(*20*), motivating complementary systems that can resolve calcium dynamics in living tissue at early developmental stages(*21*). The nematode *Caenorhabditis elegans* provides a complementary system with key advantages for dissecting calcium regulation *in vivo*. It has a long history as a Duchenne model, beginning with *dys-1* mutants and subsequent work establishing conserved roles for the dystrophin glycoprotein complex in muscle function and pathology, enabling modifier genetics and drug testing(*22*). Its transparency, short life cycle, and genetic tractability make it well suited to resolve calcium dynamics at early developmental stages using live imaging and targeted gene perturbation(*23*). Perturbations of excitability and calcium entry pathways can strongly modulate dystrophic outcomes, providing a practical way to separate contractile load effects from primary defects in calcium handling(*24*). Importantly, prior work provided evidence of muscle calcium dysregulation early in development, before overt structural damage is detectable.

Here, we test two linked hypotheses: first, that the early calcium dysregulation and the subsequent elevation in baseline calcium is present at pre-contractile developmental stages, and second, that reduced ATPase-mediated calcium extrusion across the plasma membrane contributes to this dysregulation. We combine *in vivo* calcium imaging and genetic perturbations in *C. elegans* to probe the contributions of the PMCA homologs *mca-1* and *mca-2* to plasma membrane calcium export in a dystrophic context. We validate our findings through measurements and perturbations of PMCA in patient-derived human myogenic cultures(*25–27*). These cells enable detailed assessment of calcium handling in human dystrophic muscle and allow experimental manipulation of gene expression and pharmacological modulation(*28*, *29*). Critically, they allow us to test whether elevated calcium levels and PMCA deficits are already present at the myoblast stage, before myotube formation and contractile load, and then to connect those early differences to calcium handling under repeated electrical stimulation in differentiated myotubes. Specifically, we assessed PMCA to test whether altered extrusion tracks with early calcium elevation(*30*), including PMCA immunoreactivity, PMCA1 targeted suppression, and pharmacological inhibition. We find that a conserved limitation in PMCA-mediated calcium extrusion is a contributor to a dystrophic calcium set point shift during early development that is independent of contraction-induced calcium pathologies. While calcium abnormalities during myogenesis have been reported in other systems(*31*), our study provides the first cross-species mechanistic dissection with complementary perturbations that implicate impaired calcium extrusion as an early node in the disease process.

## Results

### Early dystrophic larvae and embryos show increased muscle calcium

To determine whether dystrophic muscle exhibits elevated cytosolic calcium early in development, we expressed the muscle-specific GCaMP2 reporter *Pmyo-3::GCaMP2::unc-54 3’UTR* in either wild type (N2) or *dys-1(eg33)* worms. Pseudocolor maps of L1 body wall muscle showed higher GCaMP2 signal in dystrophic larvae relative to healthy controls (**Fig. 1a**).

**Figure 1.**
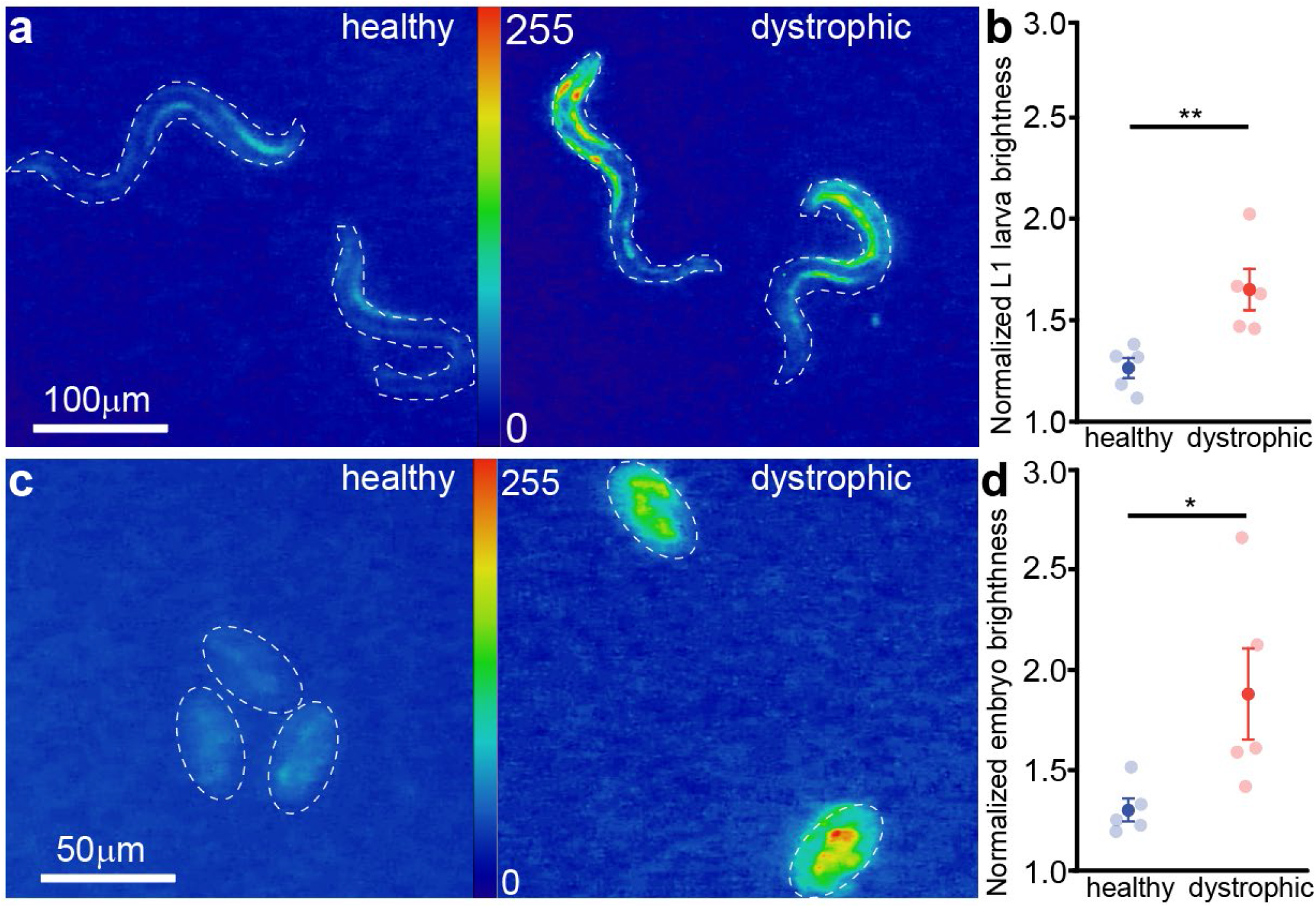
Early dystrophic worms and embryos show increased muscle calcium. **a)** Representative pseudocolor maps of body wall muscle GCaMP2 signal in healthy (N2) and dystrophic *dys-1(eg33)* L1 larvae expressing *Pmyo-3::GCaMP2::unc-54 3’UTR*, with the animal outline indicated by a dashed contour (scale bar, 100 µm; intensity scale, 0 to 255). **b)** Quantification of L1 larval brightness for healthy and dystrophic animals normalized to the background. **c)** Representative pseudocolor maps of GCaMP2 signal in three-fold stage embryos from healthy (N2) and dystrophic *dys-1(eg33)* animals (scale bar, 100 µm; intensity scale, 0 to 255), with embryos outlined by dashed contours. **d)** Quantification of embryo brightness normalized to the background. Transparent points indicate (n = 5) independent replicates, opaque points indicate group means with error bars (mean ± SEM), and significance is indicated on the plots (*p < 0.05, **p < 0.01). Statistical comparisons by Student’s unpaired t-test.

Quantification confirmed significantly higher fluorescence in *dys-1(eg33)* animals (**Fig. 1b**, healthy: 1.27 ± 0.050; dystrophic: 1.66 ± 0.103, Student’s t-test, p = 0.009), suggesting elevated sarcoplasmic calcium in dystrophic L1 larvae. This elevation in L1 larvae could stem from post-hatch locomotion-induced mechanical membrane damage, or it could reflect pre-hatch calcium dysregulation. To separate these options, we next asked whether elevated calcium is already detectable in late embryogenesis, before sustained movement begins. At the three-fold stage of development, dystrophic embryos again showed higher GCaMP2 fluorescence than healthy embryos (**Fig. 1c,d**, healthy: 1.31 ± 0.058; dystrophic: 1.88 ± 0.227, Student’s t-test, p = 0.039), suggesting calcium dysregulation was already present before post-hatch locomotion occurred.

### Dystrophic embryos hatch later than healthy controls

Having established that dystrophic embryos contain elevated muscle calcium before hatching, this finding led us to ask whether this early physiological difference is accompanied by a detectable developmental phenotype, consistent with altered muscle function. We therefore tracked hatching kinetics in synchronized embryos using timelapse imaging. Cumulative hatching data were fitted with a three-parameter logistic function of the form f = a/(1 + exp(−(x − x0)/b)), from which two parameters were extracted to characterize hatching kinetics. The first, hatching window (b), reflects the duration of the time window over which the population transitions from unhatched to hatched, where a larger value indicates a more gradual and dispersed hatching process. The second, median time to hatching (T50, x0), represents the time post-fertilization at which 50% of embryos have hatched, providing a measure of overall developmental timing. Synchronized embryos showed a significant delay in hatching rates of dystrophic embryos post-fertilization (**Fig. 2a**). Dystrophic worms showed a significantly larger hatching window than healthy controls (dystrophic: 62.3 ± 5.11 minutes post-fertilization; healthy: 44.8 ± 2.25 minutes post-fertilization, Student’s t-test, p = 0.014) indicating a prolonged and less synchronous hatching. Dystrophic embryos also showed a significantly later T50 than controls (**Fig. 2c**, dystrophic: 872 ± 24.8 minutes post-fertilization; healthy: 746 ± 10.5 minutes post-fertilization, Student’s t-test, p = 0.002), demonstrating delayed hatching in dystrophic embryos.

**Figure 2.**
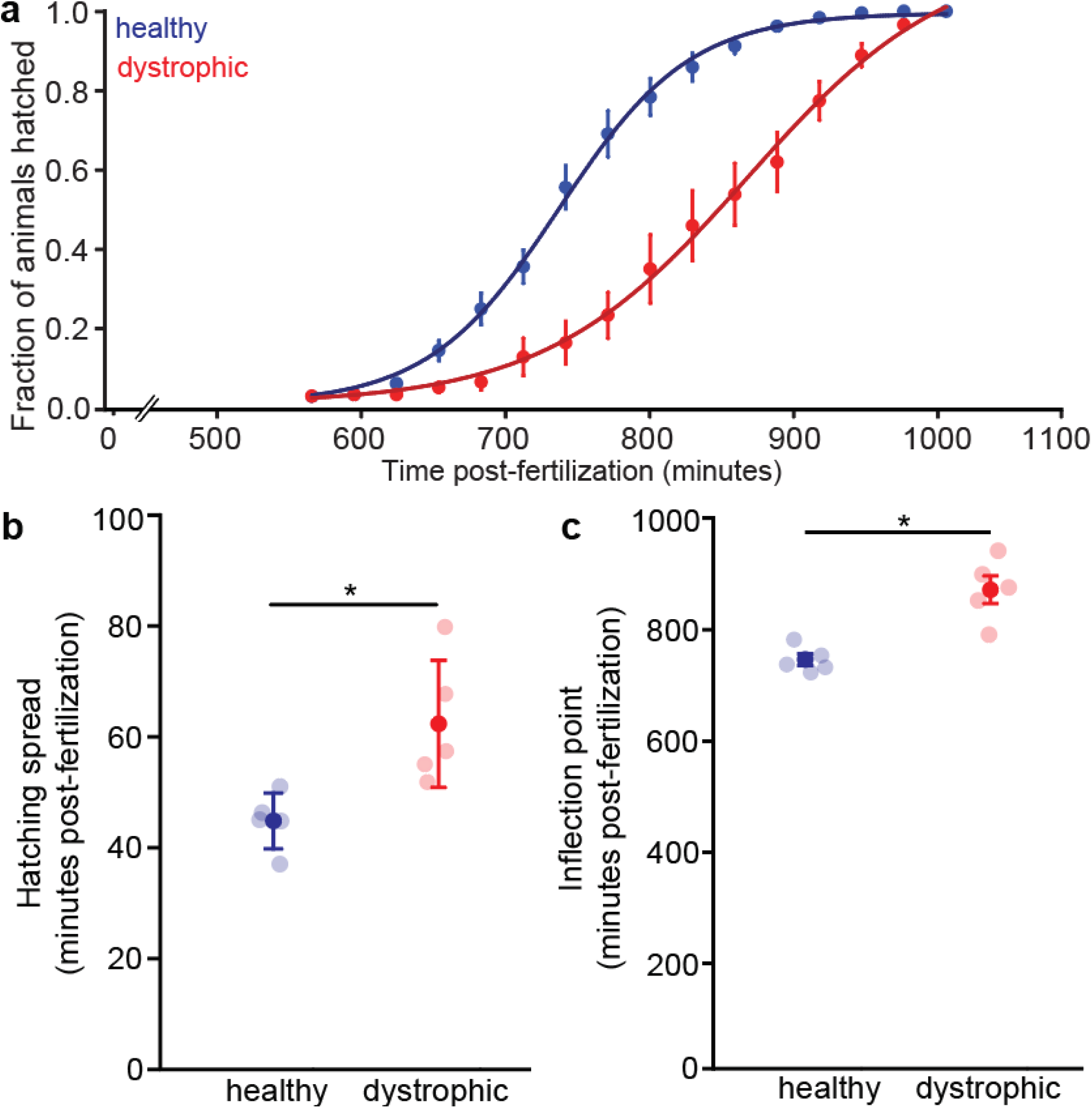
Dystrophic embryos hatch later than healthy controls. **a)** Hatching kinetics plotted as fraction hatched versus time post-fertilization for healthy (N2) and dystrophic *dys-1(eg33)* embryos, with sigmoid fits overlaid. **b)** Hatching window values extracted from the fitted hatching curves. **c)** Inflection (halfway) timepoints extracted from the fitted curves. Transparent points represent (n = 5) independent replicates, opaque points indicate group means with error bars (mean ± SEM), and significance is indicated (*p < 0.05). Statistical comparisons by Student’s unpaired t-test.

### Dystrophic adults retain more mature embryos, and eggshell weakening rescues hatching delay but does not restore normal calcium homeostasis

Delayed hatching in dystrophic animals could be a result of multiple phenomena: On the embryo side, slower developmental progression or an already existing muscle weakness before hatching could contribute to this phenotype. In the latter scenario, the embryo would show regular development, but struggle to break through the eggshell as this requires substantial muscle force production. On the parental side, longer egg retention *in utero* due to muscle weakness of the gravid adult could also delay hatching. To separate these possibilities, we asked whether dystrophic embryos progress through embryonic stages at a normal rate, whether they are retained more in gravid adults, and then tested whether weakening the eggshell would normalize the calcium elevation and allow the embryo to hatch at the normal rate.

To test whether delayed hatching reflects slower progression to late embryonic stages, we synchronized embryos and quantified when embryos reached the final stage before hatching (called ‘pretzel stage’). We found no significant difference between dystrophic and healthy embryos at any observed timepoint during embryogenesis (**Fig. 3a**), suggesting that dystrophic eggs develop normally and reach the final developmental stage at the same time as healthy eggs. To address whether eggs are retained more in gravid dystrophic adults, we examined embryos retained *in utero* and what developmental stage they are at (**Fig. 3b**). In healthy adults, almost all eggs were in the gastrula stage, and almost no eggs were found at later developmental stages. This aligns with previous findings that eggs only develop to later stages (blastula, comma, twofold, twitch and pretzel) after being laid. In contrast, in dystrophic adults, fewer eggs were at the gastrula stage (dystrophic: 0.836 ± 0.027; healthy: 0.94 ± 0.005; Welch’s t-test, p = 0.027), while significantly more retained eggs were found to be at the pretzel stage (**Fig. 3c**, Mann-Whitney U: 3969, p<0.001). This supports the idea that eggs are retained for longer and beyond the normal developmental stages in dystrophic adults.

**Figure 3.**
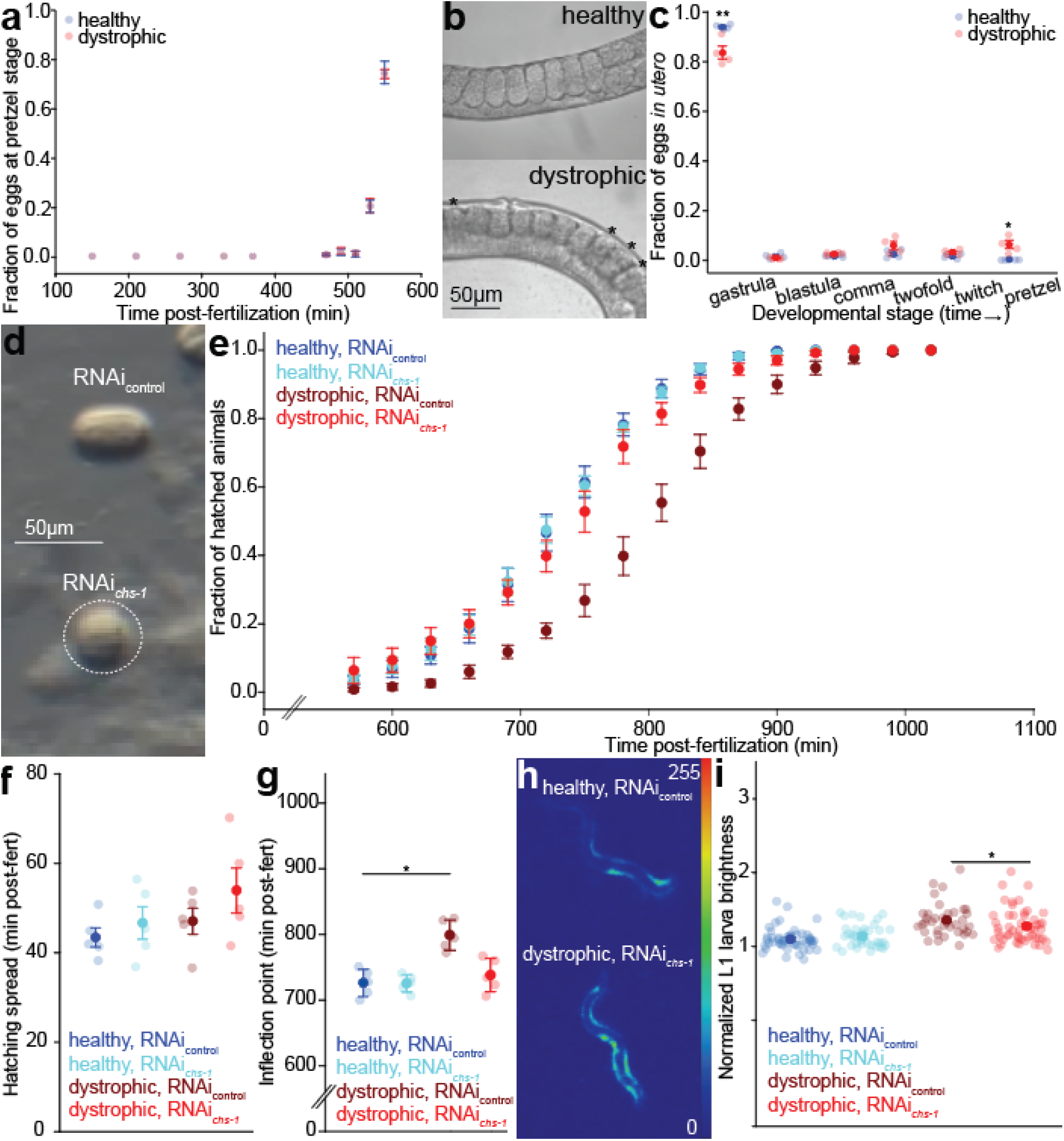
Reducing eggshell stiffness rescues dystrophic hatching delay, while post-hatch larvae retain elevated muscle calcium. **a)** Embryo staging time course showing that healthy (N2) and dystrophic *dys-1(eg33)* embryos reach the pretzel (threefold) stage with similar timing, plotted as the pretzel-staged embryo ratio versus time post-fertilization. Points represent group means of independent replicates with error bars (mean ± SEM). n = 3 independent replicates of 39-50 embryos per group. **b)** Representative images of healthy (N2) and dystrophic *dys-1(eg33)* adults illustrating differences in egg retention, with retained embryos visible in utero (scale bars, 50 µm). Asterisks represent embryos retained into later developmental stages in gravid adult dystrophic worms **c)** Distribution of developmental stages of embryos retained gravid adults, showing genotype-dependent differences in the fraction of embryos at specific embryonic stages (gastrula through threefold). Transparent points represent independent replicates, opaque points indicate group means with error bars (mean ± SEM). n = 4 independent replicates of 15-28 embryos per group. Statistical comparisons by Student’s unpaired t-test (gastrula fraction) and Mann–Whitney U test (pretzel fraction). d) Representative images of embryos after control (L4440) RNAi or *chs-1* RNAi, showing altered eggshell morphology in *chs-1* RNAi-treated embryos (dashed outline, scale bars, 50 µm). **e)** Hatching curves for healthy (N2) and dystrophic *dys-1(eg33)* embryos under L4440 control or *chs-1* RNAi conditions, showing rescue of dystrophic hatching kinetics by *chs-1* RNAi. Points indicate group means of independent replicates with error bars (mean ± SEM). n = 5 independent replicates of 63-164 embryos per condition **f, g)** Hatching window, <u>defined as the duration of the hatching window over which the majority of embryos transition from unhatched to hatched</u> (F) and T50, defined as the timepoint in which half of all embryos have hatched (G) parameters extracted from sigmoid fits to the hatching curves in (E); Transparent points represent independent replicates opaque points indicate group means with error bars (mean ± SEM). n = 5 independent replicates of 63-164 embryos per condition. Statistical comparison by Student’s unpaired t-test. **h)** Representative pseudocolor maps of GCaMP2 signal in L1 larvae under L4440 control or *chs-1* RNAi conditions in healthy and dystrophic animals (intensity scale, 0 to 255). **i)** Quantification of normalized L1 larval brightness normalized to the background, showing that *chs-1* RNAi reduces dystrophic brightness relative to dystrophic control, but larvae remain elevated relative to healthy (N2). Transparent points represent individual animals; opaque points indicate group means with error bars (mean ± SEM). N = 50 (healthy L4440), 38 (healthy *chs-1* RNAi), 88 (dystrophic L4440) and 96 (dystrophic *chs-1* RNAi) animals; the animal is the test unit. Significance is indicated in the panels (* = p < 0.05 ** = p < 0.01). Lines below asterisks represent conditions being compared.

We then weakened eggshell stiffness by reducing eggshell chitin synthesis via *chs-1* RNAi and measured hatching dynamics in laid eggs. The oval shape of healthy eggs is maintained by the stiff eggshell, and when weakened is expected to become more rounded and easier to break through by the embryo during hatching. The photo in **Fig. 3d** shows that the shape of eggs indeed changed from oval to round, demonstrating that *chs-1* RNAi treatment was successful. In addition, hatching occurred earlier in dystrophic embryos. The hatching curve was significantly advanced (**Fig. 3e**), with hatching window not significantly affected by *chs-1* RNAi in either genotype (**Fig. 3f**), but a significantly earlier T50 in dystrophic *chs-1* RNAi treated embryos (**Fig. 3g**, dystrophic L4440: 799 ± 10.3 minutes post-fertilization; dystrophic chs-1 RNAi: 738 ± 11.4, Student’s t-test, p = 0.0041). In contrast, there was no effect on healthy embryos, suggesting that eggshell barrier effects contribute to delayed hatching selectively in dystrophic embryos.

Importantly, *chs-1* RNAi did not eliminate the early calcium elevation in dystrophic embryos. Post-hatch dystrophic L1 larvae maintained significantly increased calcium levels when compared to all other groups (**Fig. 3h**: pseudocolored photos; **Fig. 3i**: Kruskal-Wallis one-way ANOVA on ranks, H = 64.8, df = 3, p < 0.001). Dunn’s post hoc test revealed that dystrophic larvae remained significantly brighter than both healthy controls (median: 1.31, IQR 1.21–1.43 vs. 1.07, IQR 1.03–1.15, p < 0.001) and healthy *chs-1* RNAi animals (p < 0.001). In addition, dystrophic *chs-1* RNAi larvae were also significantly brighter than healthy controls (median: 1.22, IQR 1.12–1.36, p < 0.001) and healthy *chs-1* RNAi animals (p = 0.022). Together, these data indicate that eggshell weakening reduced the dystrophic post-hatch calcium elevation but did not eliminate it completely. The fluorescence of healthy *chs-1* RNAi larvae did not differ from healthy controls (p = 1.000), confirming that *chs-1* RNAi did not by itself alter basal calcium in healthy animals. These results suggest that the mechanical struggle to break through the eggshell contributes to delayed hatching but cannot fully account for the calcium elevation observed in dystrophic larvae, pointing to an intrinsic difference in calcium handling.

### Embryonic calcium elevation is not fully explained by movement

The previous experiments led us to hypothesize that a locomotion-independent mechanism intrinsic to the dystrophic embryo contributes to the elevated calcium set point. To test this, we examined calcium dynamics over the entire course of embryogenesis, asking whether calcium in dystrophic embryos is already abnormally heightened before movement begins. Plotting normalized embryo brightness across development revealed that dystrophic embryos display higher fluorescence than healthy embryos throughout most of embryogenesis (**Fig. 4a**). At later developmental stages, when movement is present in the form of twitching, elevated calcium levels are consistent with activity-dependent calcium entry and contraction-induced damage to the sarcolemma. However, calcium levels were already increased 400 minutes post-fertilization, before movement became detectable in our assay at approximately 480 minutes post-fertilization, and before the first twitches reported for this species at approximately 430 minutes(*32*, *33*) (healthy: 1.008 ± 0.0185; dystrophic: 1.121 ± 0.010, Student’s unpaired t-test, t = -5.380, df = 8, p = 0.00066; all five dystrophic replicate means exceeded all five healthy replicate means). Calcium elevation at these precontractile timepoints must be movement-independent, identifying an intrinsic component of the dystrophic calcium elevation that establishes before muscle contraction begins.

**Figure 4.**
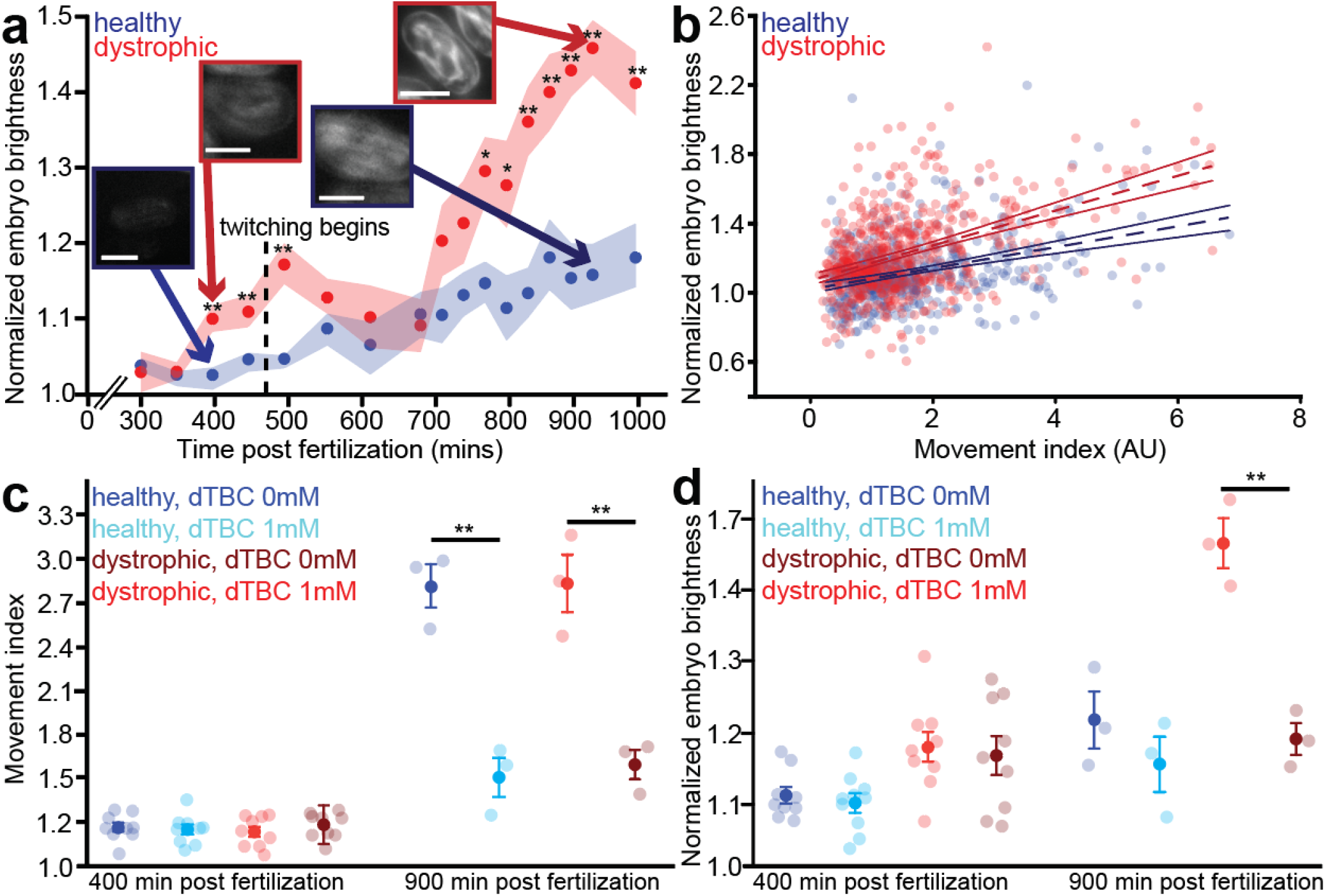
Elevated muscle calcium in dystrophic embryos is only partly explained by movement. **a)** Time course of normalized embryo brightness normalized to the background versus time post-fertilization for healthy (N2) and dystrophic *dys-1(eg33)* embryos. The dashed vertical line marks the onset of twitching. Opaque points represent group means of independent replicates (n = 5 independent replicates per group, except n = 10 at 450 and 500 mpf), shaded transparent region represents error (mean ± SEM). Representative embryo images at selected timepoints are shown. n = 5 independent replicates of 20 embryos each per genotype, scale bars, 25 µm. **b)** Normalized brightness plotted as a function of movement index for individual embryos, with linear fits shown for each genotype (healthy: R^2^ = 0.122, standard error of estimate (SEE) = 0.174, dystrophic R^2^ = 0.204, SEE = 0.221. N = 538 embryos per genotype **c)** Movement index at approximately 400 and 900 minutes post-fertilization for embryos treated with water (0 mM dTBC) or 1 mM d-tubocurarine chloride (dTBC). n = 9 independent replicates of 20 embryos per group at 400 mpf; n = 3 independent replicates of 20 embryos per group at 900 mpf **d)** Normalized embryo brightness at the same timepoints and conditions as (C) with the same sample sizes. Transparent points represent independent replicates, opaque points indicate group means with error bars (mean ± SEM). n = 9 independent replicates of 20 embryos per group at 400 mpf; n = 3 independent replicates of 20 embryos per group at 900 mpf. Statistical comparisons in (C) and (D) by Mann–Whitney U test or Student’s unpaired t-test as appropriate; significance is indicated on the plots (*p < 0.05, **p < 0.01).

Although calcium elevation precedes the onset of movement, we considered that once movement occurs, there could be differences in the amount of movement between controls and dystrophic embryos that could account for the higher calcium levels in dystrophic embryos. To quantify movement, we calculated a normalized pixel variance index along the longitudinal axis of each embryo (see Methods), with a higher index indicating more movement. We then plotted fluorescence as a function of the movement index for individual embryos across both genotypes (**Fig. 4b**). A linear regression showed that dystrophic embryos were brighter than healthy embryos even at the lowest levels of movement: the y-intercept was significantly higher than in healthy embryos (dystrophic = 1.08 ± 0.016; healthy = 1.03 ± 0.013 F(1,1072) = 5.06, p = 0.0246). Additionally, dystrophic embryos displayed a significantly greater increase in fluorescence with movement, indicated by a significant genotype × movement interaction (slope: dystrophic = 0.1 ± 0.008; healthy = 0.0594 ± 0.006, p = .00028, ANCOVA, n=538 embryos per genotype), indicating that movement drives a proportionally greater calcium increase in dystrophic embryos. Regression fits of fluorescence intensity (y) vs movement index (x) also only explain a small portion of the variance in both genotypes (healthy R^2^ = 0.122; dystrophic R^2^ = 0.204), suggesting that movement is not the only predictor of fluorescence differences between healthy and dystrophic embryos. Together, these findings suggest that dystrophic muscle already begins with a higher calcium baseline and responds with a larger calcium increase to mechanical activity.

The findings in Figs 4A and 4B suggest that the dystrophic elevation in calcium cannot be fully explained by movement. To further test whether movement alone drives calcium elevation, we pharmacologically reduced embryonic movement using d-tubocurarine chloride (dTBC) in gravid adults. dTBC is a competitive antagonist of nicotinic acetylcholine receptors and blocks motoneuronal innervation of skeletal muscles(*34*). We reasoned that if movement were driving calcium elevation, suppressing it by blocking neuromuscular innervation should eliminate or substantially reduce the calcium level differences between dystrophic and healthy embryos.

We first quantified embryonic movement across genotypes. Embryos were harvested after cessation of movement was observed in gravid adults, i.e., after the drug’s effect was established, and monitored at key timepoints of embryogenesis. At 400 minutes post-fertilization, more than one hour prior to the expected onset of robust embryonic movement, no movement was observed and movement indices did not differ significantly between dTBC-treated and untreated embryos within either genotype (**Fig. 4c**, healthy 0mM dTBC: 1.17 ± 0.032; healthy 1mM dTBC: 1.16 ± 0.034, Student’s t-test, t = 0.349, df = 16, p = 0.732; dystrophic 0mM dTBC: 1.14 ± 0.035; dystrophic 1mM dTBC: 1.18 ± 0.030, p = 0.350). This demonstrates that blocking motoneuronal innervation had no effect on embryonic movement at this developmental stage and strongly suggests that muscle activation was absent in all groups. At 900 minutes post-fertilization (about 7 hours after twitching had commenced), blocking motoneuronal input significantly reduced movement within each genotype (healthy: Mann-Whitney rank-sum U = 0.000; dystrophic: Student’s t-test, t = 5.614, df = 4, p = 0.00495), indicating that movement was successfully dampened by the drug in both healthy and dystrophic embryos. Despite this reduction, movement at 900 mpf did not differ significantly between genotypes, either untreated (dystrophic: 2.833 ± 0.196; healthy: 2.817 ± 0.148, p = 0.95) or dTBC-treated (dystrophic: 1.597 ± 0.100; healthy: 1.512 ± 0.133, p = 0.64), indicating that dTBC reduced movement comparably in both genotypes.

Having confirmed that dTBC successfully reduced movement at 900 minutes post-fertilization, we next asked whether this reduction in movement was accompanied by a reduction in calcium levels. Because dTBC was delivered maternally and embryos were rinsed and imaged on drug-free plates afterward, movement suppression relied on dTBC retained within the embryos after parental exposure rather than on continuous exposure during imaging. Retained drug reduced, but did not abolish, movement at 900 minutes post-fertilization, since treated embryos remained above the movement index floor of immobile embryos. At 400 minutes post-fertilization no movement was detectable in any group. That absence rests on the movement index and on the developmental absence of functional neuromuscular junctions, not on the drug. Under these conditions, dystrophic embryos showed significantly higher calcium levels than healthy embryos at 900 minutes post-fertilization in the absence of dTBC (**Fig. 4d**). At 400 minutes post-fertilization (mpf), when movement had not yet begun, calcium was already significantly elevated in dystrophic embryos in comparison to healthy embryos, as established above (**Fig. 4a**). At 900 mpf, dTBC treatment significantly reduced calcium levels in dystrophic embryos (1.463 ± 0.0347 to 1.191 ± 0.0224, Student’s t-test, p = 0.00274; Welch’s t-test, p = 0.00467, applied because the Brown-Forsythe test indicated unequal variances) but did not significantly affect healthy embryos (1.217 to 1.155, p = 0.326). Following dTBC treatment, calcium levels in dystrophic embryos became statistically indistinguishable from those in untreated healthy embryos (p = 0.60), indicating that reducing movement substantially, though not completely, normalized the dystrophic calcium elevation. At 400 mpf, dTBC had no effect on calcium in either genotype (healthy: 1.100 ± 0.010 untreated versus 1.102 ± 0.014 dTBC, p = 0.90; dystrophic: 1.180 ± 0.021 versus 1.168 ± 0.026, p = 0.75; Student’s unpaired t-test, n = 9 replicates of 20 embryos per group), and dystrophic embryos remained brighter than healthy embryos in the presence of the drug (1.168 ± 0.026 versus 1.102 ± 0.014, difference 0.067, 95% CI 0.004 to 0.130, Student’s t-test, p = 0.039; Welch’s t-test, p = 0.044). The dystrophic offset is therefore present with cholinergic drive blocked and before movement begins. These data are consistent with an elevated calcium baseline level in dystrophic embryos that can be modulated by movement but is not created by movement alone.

### Targeted RNAi perturbations identify calcium export limitation and implicate plasma membrane calcium ATPases

Having established that the dystrophic calcium elevation cannot be explained by movement alone, we reasoned that an intrinsic defect in calcium handling was likely contributing to the elevated calcium baseline. To identify which calcium regulatory processes might contribute, we performed a targeted RNAi screen spanning pathways expected to alter calcium levels through different routes. The first potential route targets sarcomeric contractility and membrane excitability. Despite our embryo data arguing against movement being the sole driver of elevated calcium, contractile activity drives calcium entry through excitation-contraction coupling. Prior work from our lab demonstrated that targeted RNAi silencing of calcium handling genes including calmodulin, SERCA, and calsequestrin produces distinct and interpretable effects on basal calcium levels in dystrophic *C. elegans*, validating this approach for understanding calcium regulatory pathways in vivo(*24*). Genes examined in that earlier screen appear in red in Fig. 5a and were not retested here: *sca-1* knockdown raised basal calcium in both genotypes, *cmd-1* knockdown returned dystrophic calcium to healthy levels, and *csq-1* knockdown altered calcium transient amplitude in opposite directions in the two genotypes. The ryanodine receptor unc-68 could not be included, as the *C. elegans* RNAi feeding library contains no clone targeting it. Building on this foundation, we expanded the RNAi screen to include sarcomeric contractility genes, voltage-gated calcium channels, and plasma membrane calcium ATPases, reasoning that each category would alter cytosolic calcium through a mechanistically distinct route (**Fig. 5a**). The first group included contractility and excitability genes to quantify how much of the adult dystrophic calcium elevation is activity-dependent, and to provide a baseline to interpret the calcium export results. The second group targeted voltage-gated ion channels, which mediate depolarization-driven entry and represent a direct route by which excitability changes could alter cytosolic calcium load independently of mechanical force. The third group targeted plasma membrane calcium ATPases, which export calcium against its electrochemical gradient and, together with sodium calcium exchange, maintain low resting cytosolic calcium independently of contractile activity. Sodium calcium exchangers were not manipulated here, so these experiments address the ATPase contribution rather than plasma membrane export in total. We reasoned that if a limitation in calcium export contributes to the elevated dystrophic set point, knockdown of export genes should produce a different calcium response in dystrophic versus healthy animals compared to the activity-dependent perturbations. These experiments were carried out in adult animals. Across panels, healthy controls and dystrophic animals differed significantly in both swimming frequency and muscle fluorescence (Student’s t-test, p < 0.001 for all comparisons), confirming the baseline differences between genotypes. For each measure and genotype, we fit a single one-way ANOVA across all nine RNAi conditions (L4440 control plus *lev-11*, *myo-3*, *pat-10*, *unc-38*, *egl-19*, *unc-2*, *mca-1*, and *mca-2*; n = 4 per group), with Holm-Šídak post hoc comparisons made against the shared L4440 control (brightness, healthy: F(8,27) = 18.844; brightness, dystrophic: F(8,27) = 13.200; swimming, healthy: F(8,27) = 86.098; swimming, dystrophic: F(8,27) = 22.450)

**Figure 5.**
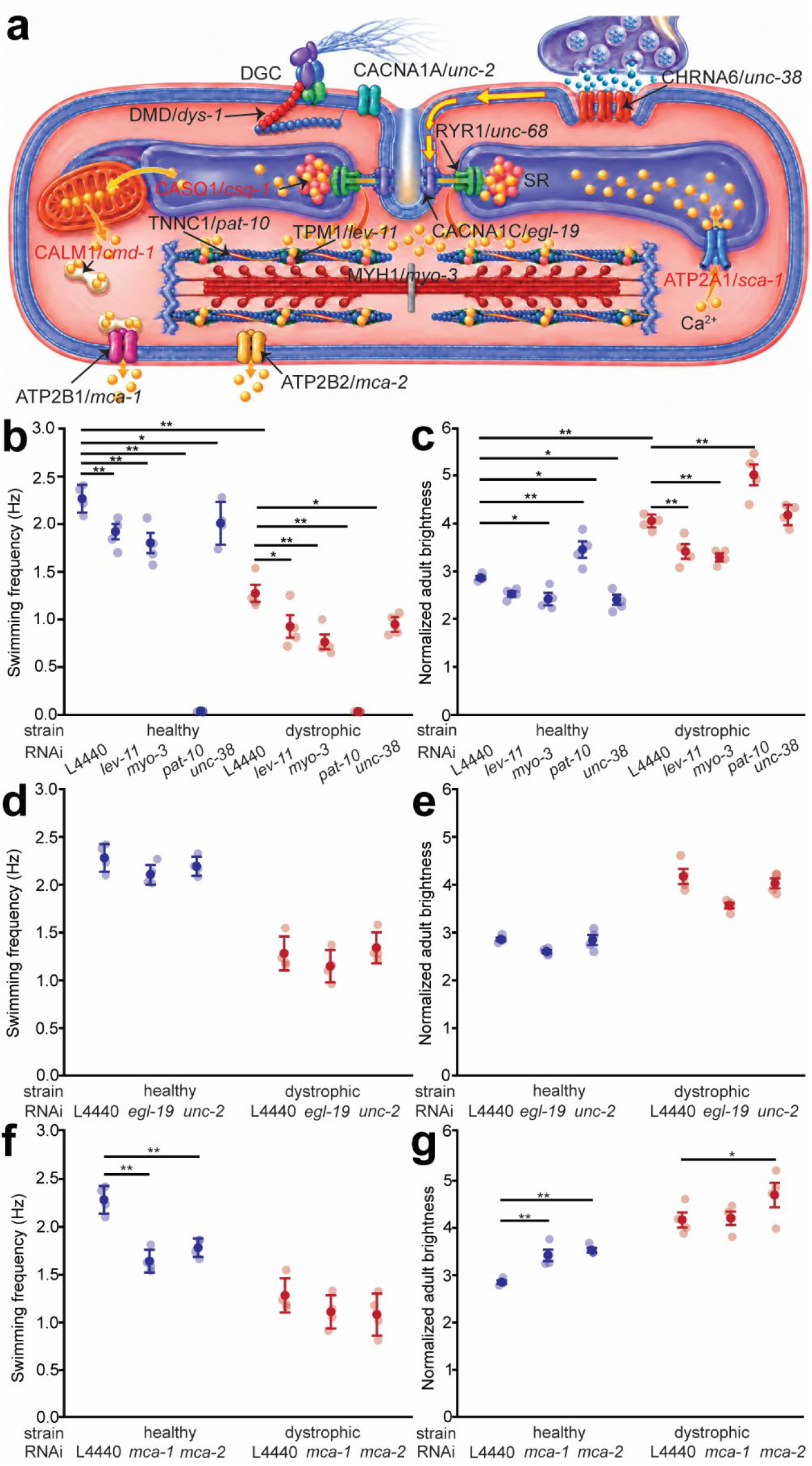
RNAi perturbations of contractility, excitability, and calcium transport identify mca-1 as a limiting calcium extrusion pathway. **a)** Schematic of the RNAi screen logic. Targeted genes are grouped by expected effect on cytosolic calcium: muscle contractility (*lev-11*, *myo-3*, *pat-10*) and excitability (*unc-38*, *egl-19*, *unc-2*) contribute to muscle activity, while plasma membrane calcium ATPases (*mca-1*, *mca-2*) mediate export. *mca-1* is the diagnostic test of export reserve; *mca-2* serves as a dynamic-range control. Labels in red (*cmd-1*, *csq-1*, *sca-1*) denote calcium handling genes examined in our earlier screen(*24*) and shown here for context rather than retested. The ryanodine receptor *unc-68* is shown for completeness; no clone targeting it is present in the *C. elegans* RNAi feeding library, so it could not be included in a feeding screen. Remaining labelled components (DGC, *dys-1*, SR) indicate anatomical context. Conceptual schematic; no data shown. **b)** Swimming frequency (Hz) for wild-type (WT) and dystrophic (*dys-1*) animals following RNAi targeting *lev-11*, *myo-3*, *pat-10*, or *unc-38.* (L4440, RNAi control). **c)** Brightness normalized to the background under the same conditions as (b). **d)** Swimming frequency after *egl-19* or *unc-2* RNAi. **e)** Normalized muscle fluorescence after *egl-19* or *unc-2* RNAi. **f)** Swimming frequency after *mca-1* or *mca-2* RNAi. **g)** Normalized muscle fluorescence after *mca-1* or *mca-2* RNAi. *mca-1* RNAi elevates fluorescence in WT toward dystrophic levels with only a modest additional effect in *dys-1*, whereas *mca-2* RNAi increases fluorescence in both genotypes, indicating that dystrophic animals are not at a hard fluorescence ceiling. Transparent points represent independent replicates, opaque points indicate group means with error bars (mean ± SEM). n = 4 independent replicates of 10 worms each per condition Data for RNAi control (L4440) is the same dataset serving each swimming and brightness panel for both genotypes. Statistical comparisons by one-way ANOVA with Holm-Šídák post hoc correction for normally distributed data; significance indicated on the panels (*p < 0.05, **p < 0.01).

Knockdown of sarcomeric contractility genes reduced swimming frequency in healthy and dystrophic animals (**Fig. 5b**, one-way ANOVA with Holm-Šídák post hoc, p < 0.001). In healthy animals, *lev-11* (2.28 ± 0.072 Hz vs. 1.93 ± 0.082 Hz, p = 0.006), *myo-3* (1.81 ± 0.107 Hz, p = 0.004), and *unc-38* RNAi (1.91 ± 0.112 Hz, p = 0.014) significantly reduced swimming relative to healthy controls (2.28 ± 0.072 Hz), and *pat-10* RNAi caused full paralysis (0.003 ± 0.003 Hz, p < 0.001). Consistent with movement-induced calcium load contributing to the dystrophic phenotype, *lev-11* and *myo-3* knockdown reduced muscle calcium fluorescence in dystrophic animals **(Fig. 5c**, one-way ANOVA with Holm-Šídák post hoc, omnibus p < 0.001; *lev-11*: 3.42 ± 0.150, p = 0.023; *myo-3*: 3.31 ± 0.058, p = 0.011, relative to dystrophic control animals: 4.17 ± 0.158). In contrast, *pat-10* knockdown significantly increased muscle calcium fluorescence relative to controls in both genotypes (dystrophic: 5.02 ± 0.222, p = 0.0114; healthy: 3.46 ± 0.17, p = 0.0022), consistent with troponin C acting as a cytosolic calcium buffer, such that its loss raises free calcium even as it abolishes movement.

Similarly, knockdown of voltage-gated calcium channels reduced swimming in both genotypes (**Fig. 5d**, one-way ANOVA with Holm-Šídák post hoc, omnibus p < 0.001).In healthy animals, *egl-19* RNAi did not significantly alter swimming (1.93 ± 0.112 Hz, p = 0.188 vs. healthy controls 2.28 ± 0.072 Hz) or muscle calcium fluorescence (Fig. 5e, 2.60 ± 0.033, p = 0.165 vs. healthy controls 2.86 ± 0.036), while *unc-2* RNAi did not significantly alter fluorescence (2.84 ± 0.106, p = 0.918) relative to controls. While silencing of contractility and excitability genes confirmed that movement-dependent calcium load contributes to the dystrophic phenotype in adult animals, these perturbations cannot fully account for the calcium elevation observed in precontractile embryos where neither sarcomeric contractility nor excitation-contraction coupling are operating yet. Increased membrane excitability could in principle drive calcium entry through voltage-gated channels irrespective of movement.

However, at embryonic stages, before the excitation-contraction coupling apparatus has matured, a sustained elevation in resting calcium is more likely to reflect a defect in calcium clearance than in calcium entry alone. We therefore targeted the plasma membrane calcium ATPases *mca-1* and *mca-2*, the primary calcium export pathway at the plasma membrane, to test whether impaired extrusion could account for the movement-independent component of the dystrophic calcium set point shift. Although *mca-1* and *mca-2* RNAi both reduced swimming frequency in healthy animals (**Fig 5f**, one-way ANOVA with Holm-Šídák post hoc, omnibus p < 0.001; *mca-1*: 1.64 ± 0.060 Hz, p < 0.001 vs. healthy controls; *mca-2*: 1.78 ± 0.049 Hz, p < 0.001 vs. healthy controls 2.28 ± 0.072 Hz), swimming frequency was unaffected in dystrophic animals following either knockdown (one-way ANOVA, omnibus p = 0.34). Dystrophic animals treated with *mca-1* RNAi continued to swim at frequencies comparable to healthy animals given the same treatment. This, combined with other RNAi lines showing even lower swimming frequencies in dystrophic animals confirms that movement had not reached a floor, and that the absence of an effect in dystrophic animals reflects a genuine insensitivity to mca perturbation rather than an inability to detect further reductions in an already immobile animal. Critically, *mca-1* RNAi increased calcium fluorescence in healthy animals (**Fig. 5g**; 3.42 ± 0.120 vs. healthy controls 2.86 ± 0.036, Holm-Šídák p = 0.0032) while having no detectable effect in dystrophic animals (4.21 ± 0.142 vs. dystrophic controls 4.17 ± 0.158; difference of means = +0.034, 95% CI −0.49 to +0.55). In contrast, *mca-2* RNAi elevated fluorescence significantly in both healthy (3.44 ± 0.116, Holm-Šídák p = 0.0027) and dystrophic animals (5.06 ± 0.353, Holm-Šídák p = 0.0093; difference of means = 0.890 relative to dystrophic controls), demonstrating that the bounded null observed for *mca-1* in dystrophic animals was not due to a ceiling effect but instead reflects a pathway-specific limitation, since *mca-2* knockdown produced a substantial shift in the same animals on the same fluorescence axis. Taken together, these results show that reducing either *mca-1* or *mca-2* impairs calcium homeostasis and locomotion in healthy animals. In dystrophic animals, however, reducing either gene had no effect on locomotion, and only *mca-2* RNAi further impaired calcium homeostasis, whereas *mca-1* RNAi had no detectable effect. The selective absence of an *mca-1* RNAi phenotype in dystrophic animals suggests that *mca-1* function is already compromised in the dystrophic background.

### RNA sequencing provides transcriptomic context in human myogenic cultures and highlights activity-linked programs relevant to calcium handling

The above data identified an impaired *mca-1-*mediated calcium export as an underlying factor in elevating calcium in dystrophic muscle and implicated this deficit to be present before muscle movement occurs. If this impairment was conserved in human dystrophic muscle, this would implicate DMD as a developmental disorder. We thus tested whether a homolog impairment exists in human muscle, whether it is already present before differentiation of myoblasts into contractile muscle fibers, and whether defective calcium cycling persists in myotubes under electrical stimulation in the absence of mechanical load. Patient-derived human healthy and dystrophic myoblasts offer a tractable system for this test, as calcium dynamics can be assessed before and during myotube formation as well as before the mature contractile stage. Before performing targeted perturbations of *PMCA,* the human homolog of the *mca* family in *C. elegans*, we first sought to establish whether dystrophic myoblasts show broad transcriptomic differences consistent with altered calcium handling, both to validate the system and to provide pathway-level context for the mechanistic experiments that follow. We therefore profiled transcriptomes from healthy and dystrophic human myogenic cultures (three independent populations per condition). Principal component analysis showed robust genotype-based segregation of altered expression (**Fig. 6a)**, and differential expression identified 2137 genes with significant changes in dystrophic myoblasts (**Fig. 6b**; adjusted p < 0.05).

**Figure 6.**
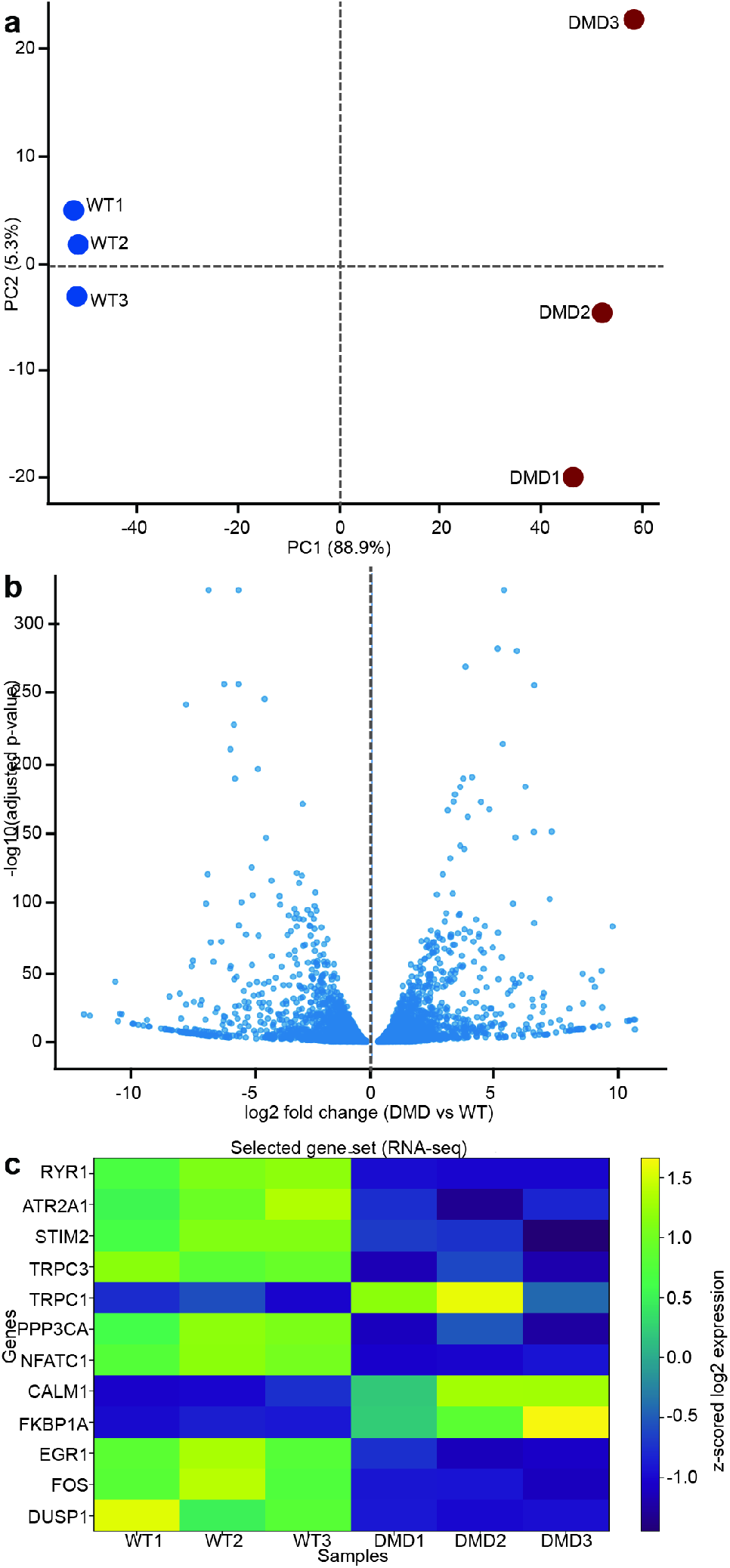
RNA sequencing validates separation of WT and dystrophic human myogenic cultures and reveals coordinated changes in calcium handling and activity-dependent transcriptional programs. **a)** Principal component analysis of transcriptome profiles from WT (WT1 to WT3) and DMD (DMD1 to DMD3) cultures, showing robust genotype-based segregation. **b)** Volcano plot summarizing differential expression (DMD versus WT) across detected transcripts, plotted as log2 fold change versus −log10 adjusted p-value. Differential expression was assessed by the Wald test within the DESeq2 framework with Benjamini–Hochberg correction; significance threshold: adjusted p < 0.05. **c)** Heatmap of a curated gene set relevant to calcium homeostasis and downstream signaling (RYR1, ATP2A1, STIM2, TRPC1, TRPC3, PPP3CA, NFATC1, CALM1, FKBP1A, EGR1, FOS, DUSP1), displayed as row z-scored log2 expression across samples. These transcriptome-level differences support the use of this culture system for mechanistic testing of PMCA-dependent calcium extrusion in subsequent experiments. WT corresponds to the healthy AB1190 cell line, and DMD corresponds to the AB1071 dystrophic cell line.

A curated gene set view (**Fig. 6c**) showed coordinated differences in calcium homeostasis and downstream signaling programs (for example RYR1, ATP2A1, STIM2, TRPC1, TRPC3, PPP3CA, NFATC1, CALM1, FKBP1A, EGR1, FOS, DUSP1), indicating that dystrophic myoblasts already differ broadly in calcium-handling gene expression before differentiation and muscle contractions. We observed downregulation of activity-dependent transcription factors with known positive regulatory roles in calcium-responsive and immediate early gene programs. In this dataset, *EGR1* (log2FoldChange = −0.865, padj = 3.02 × 10⁻¹³) and *FOS* (log2FoldChange = −0.846, padj = 1.54 × 10⁻⁹) were significantly downregulated in dystrophic myoblasts. EGR1 can transactivate the promoter of the *ATP2B2* ATPase (PMCA2) in neuronal and sensory cells, with paralog-selective effects(*35*), raising the testable possibility that reduced EGR1/FOS activity contributes to impaired PMCA-family expression; a direct regulatory link to *ATP2B1* (PMCA1) in muscle, however, remains to be established (**Fig. 6c**). Notably, *ATP2B1* was not identified as differentially expressed in this dataset (log2 fold change = −0.11, padj = 0.463).

### Reduced PMCA1 abundance or activity contributes to elevated basal calcium in dystrophic human myoblasts

Having established broad transcriptomic remodeling of calcium handling programs in dystrophic myoblasts, we turned to targeted functional and protein-level assessment of PMCA-dependent calcium export to overcome the limited statistical power inherent to RNA-seq. Baseline Fluo-4 AM calcium imaging supported elevated sarcoplasmic calcium in dystrophic (AB1071) compared to healthy (AB1190) myoblasts, i.e., before the occurrence of any muscle contractions (**Figs. 7a,b**, healthy: 8.06 ± 0.623 AU arbitrary units; dystrophic: 11.9 ± 1.33 AU, Welch’s t-test, p = 0.042). PMCA immunofluorescence using a polyclonal antibody that does not distinguish between PMCA1 and PMCA4, revealed that PMCA protein abundance was significantly reduced in dystrophic myoblasts (**Figs. 7c, d,** healthy: 58.4 ± 4.50 AU; dystrophic: 31.5 ± 7.11 AU, Student’s t-test, p = 0.013). These results suggested an impaired calcium transport across the sarcolemma.

**Figure 7.**
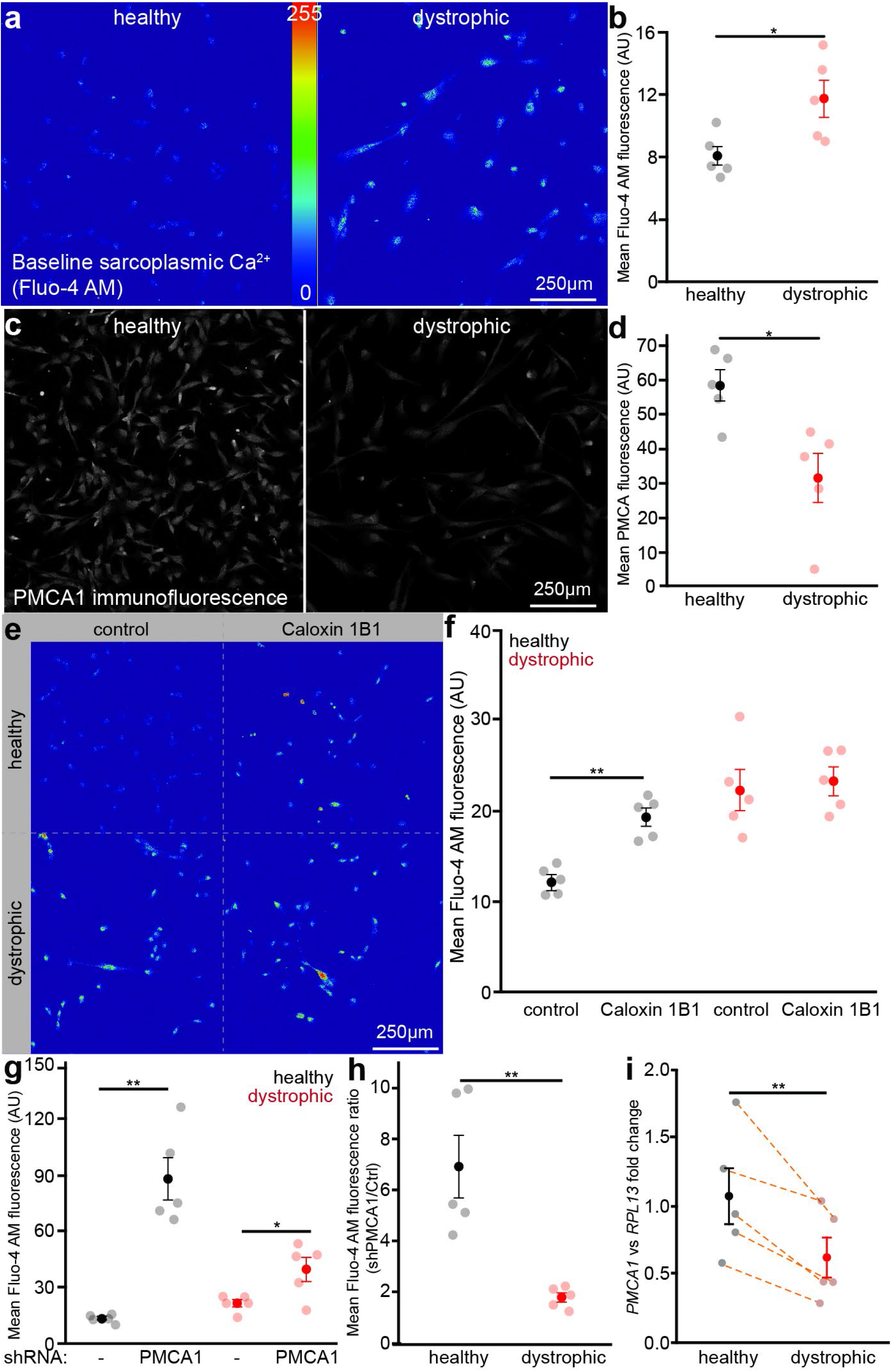
Reduced PMCA1 expression contributes to elevated basal sarcoplasmic calcium in dystrophic human myoblasts and is reproduced by genetic or pharmacological inhibition. **a)** Representative pseudocolor images of baseline sarcoplasmic Ca^2+^ (Fluo-4) in healthy (AB1190) and dystrophic (AB1071) myoblast cultures (same lookup table and intensity scale). Scale bar, 250 µm. **b)** Quantification of mean Fluo-4 AM brightness confirms elevated basal calcium in dystrophic myoblasts (Student’s unpaired t-test). n = 5 independent replicates of 111-200 cells per group. **c)** Representative anti-PMCA immunofluorescence images in healthy and dystrophic myoblasts. Scale bar, 250 µm. **d)** Quantification of mean anti-PMCA immunofluorescence shows reduced signal in dystrophic cultures (Student’s unpaired t-test). n = 5 independent replicates per group, with 83-150 cells per replicate for healthy and 77-150 for dystrophic cultures. **e)** Representative pseudocolor baseline Fluo-4 images of healthy and dystrophic myoblasts treated with water control or Caloxin 1B1. **f)** Quantification of mean Fluo-4 brightness showing Caloxin 1B1 elevates basal calcium significantly in healthy myoblasts, with no significant further increase in dystrophic cells (Student’s unpaired t-test). n = 5 independent replicates of 200 cells per group. **g)** PMCA1 silencing (PMCA1 shRNA) increases basal Fluo-4 fluorescence significantly in healthy myoblasts and produces a smaller but significant increase in dystrophic myoblasts (Student’s unpaired t-test). n = 5 independent replicates per genotype. Healthy cultures contributed 175-391 control and 17-71 PMCA1 shRNA cells per replicate; dystrophic cultures contributed 114-544 control and 8-56 PMCA1 shRNA cells per replicate. **h)** Ratio of mean Fluo-4 AM fluorescence in PMCA1 shRNA (mCherry-positive) cells relative to mCherry-negative cells from the same cultures (shPMCA1/Ctrl) within each genotype. The proportional increase is markedly greater in healthy myoblasts, consistent with a reduced reserve of PMCA1-mediated export capacity. n = 5 independent replicates as in (g), from which these ratios are computed. **i)** qPCR of PMCA1 transcripts in paired healthy and dystrophic samples normalized to *RPL13* shows reduced PMCA1 expression in dystrophic myoblasts (Student’s paired t-test); dashed lines connect matched replicates. Transparent points represent independent replicates, opaque points indicate group means with error bars (mean ± SEM). n = 5 independent replicates per group run in triplicate. Significance is indicated in the plots (*p < 0.05, **p < 0.01).

We next asked whether this potential reduction in PMCA is functionally important and sufficient to explain the concurrent calcium elevation. To test this, we pharmacologically inhibited PMCA1 and 4 activity with Caloxin 1B1 in healthy and dystrophic myoblasts, reasoning that if reduced PMCA calcium export capacity drives the dystrophic-specific calcium elevation, then acutely limiting that capacity in healthy cells should shift their calcium toward dystrophic levels but the same manipulation should not affect the already impaired dystrophic cells. Caloxin 1B1 elevated basal Fluo-4 fluorescence significantly in healthy myoblasts (**Fig. 7e, f**, control: 12.2 ± 0.668 AU; Caloxin 1B1: 19.2 ± 0.992 AU, Student’s t-test, p < 0.001) but this effect was absent in dystrophic cells (control: 22.2 ± 2.276 AU; Caloxin 1B1: 23.3 ± 1.492 AU, Student’s t-test, p = 0.704,). These findings are consistent with healthy cells having a higher PMCA-dependent calcium export capacity than dystrophic cells. Our findings in *C. elegans* had implicated *mca-1* (the PMCA1 homolog) as the limiting calcium export pathway rather than *mca-2.* Because Caloxin 1B1 inhibits both PMCA1 and PMCA4, and can thus not separate between them, we next used targeted RNAi silencing to specifically reduce PMCA1 expression. *PMCA1*-targeted silencing significantly increased basal Fluo-4 AM calcium fluorescence in healthy myoblasts (**Fig. 7g**, control: 13.8 ± 0.894 AU; PMCA1 shRNA: 88.2 ± 11.4 AU, Student’s t-test, p < 0.001). In dystrophic myoblasts, a smaller but still significant increase in calcium level was observed (dystrophic control: 21.7 ± 1.94 AU, PMCA1 shRNA: 39.8 ± 6.42 AU, Student’s t-test, p = 0.028). The ratio of calcium increase following *PMCA1* silencing was markedly greater in healthy compared to dystrophic myoblasts (**Fig. 7h**, healthy: 6.66 ± 1.18; dystrophic: 1.78 ± 0.180, Student’s t-test, p = 0.004), consistent with a reduced reserve of PMCA1-mediated calcium export capacity in dystrophic myoblasts. Together, the pharmacological and genetic loss-of-function results converge on the interpretation that PMCA1-dependent calcium export capacity is already diminished in dystrophic myoblasts, contributing to elevated calcium levels in dystrophic muscle.

To directly quantify *PMCA1* transcript levels with greater sensitivity than RNA-seq affords, we performed targeted qPCR of myoblasts. qPCR measurements detected reduced *PMCA1* transcript abundance in dystrophic samples relative to healthy controls (**Fig. 7i**, ΔΔCt difference: 0.844 ± 0.142, Student’s paired t-test, p = 0.004), consistent with the interpretation that reduced *PMCA1* expression contributes to the elevated basal calcium in dystrophic muscles.

### Dystrophic myotubes show altered calcium dynamics under repeated electrical stimulation

To determine how elevated calcium and constrained clearance affect muscles under repeated demand, such as during repeated activation, we examined differentiated, noncontractile myotubes during repeated electrical field stimulation. Myotubes were exposed to two sets of repeated trains of electrical stimuli (**Fig. 8a**; stimulus frequency: 100 Hz, interburst interval: 5 s, interval between sets: 20 s). Dystrophic myotubes began with a higher baseline fluorescence than healthy myotubes (**Fig. 8b**, healthy: 1.41 ± 0.085; dystrophic: 2.06 ± 0.133, Student’s t-test, p = 0.003) and showed a longer calcium decay time constant, indicating slower calcium clearance in dystrophic myotubes (**Fig. 8c**, τ healthy: 0.712 ± 0.085 s, τ dystrophic: 1.14 ± 0.072 s, Student’s t-test, p = 0.005). Dystrophic myotubes also significantly exceeded healthy myotubes in peak fluorescence during the first stimulation bout (**Fig. 8d**, healthy: 2.44 ± 0.131; dystrophic: 3.77 ± 0.52, Student’s t-test, p = 0.038), consistent with their elevated baseline. Peak fluorescence did not differ significantly between the first and second stimulation sets within either genotype (**Fig. 8d**, healthy set 1: 2.44 ± 0.133 AU, set 2: 2.33 ± 0.186 AU; dystrophic set 1: 3.77 ± 0.52 AU, set 2: 3.28 ± 0.523 AU), though the numerical trend toward reduced peak fluorescence in dystrophic myotubes across bouts is consistent with limited calcium recovery capacity between repeated demands. To assess whether myotubes recovered to baseline calcium levels between sets, we quantified the intermission slope, defined as the rate of change in fluorescence during the rest interval between stimulation sets. A positive slope during this interval indicates calcium is not clearing toward baseline and instead accumulating, reflecting impaired recovery. We observed a clear trend in the expected direction, as dystrophic myotubes showed a modestly positive intermission slope, and healthy myotubes showing a negative one (**Fig. 8e**, healthy: −2.90 × 10⁻⁵ ΔF/F s⁻¹ ± 5.37 × 10⁻⁶; dystrophic: 7.5 × 10⁻⁶ ± 1.42 × 10⁻⁵), consistent with impaired calcium recovery in dystrophic myotubes, though this difference did not reach significance under the Welch’s t-test, used here given unequal variances (p = 0.058)

**Figure 8.**
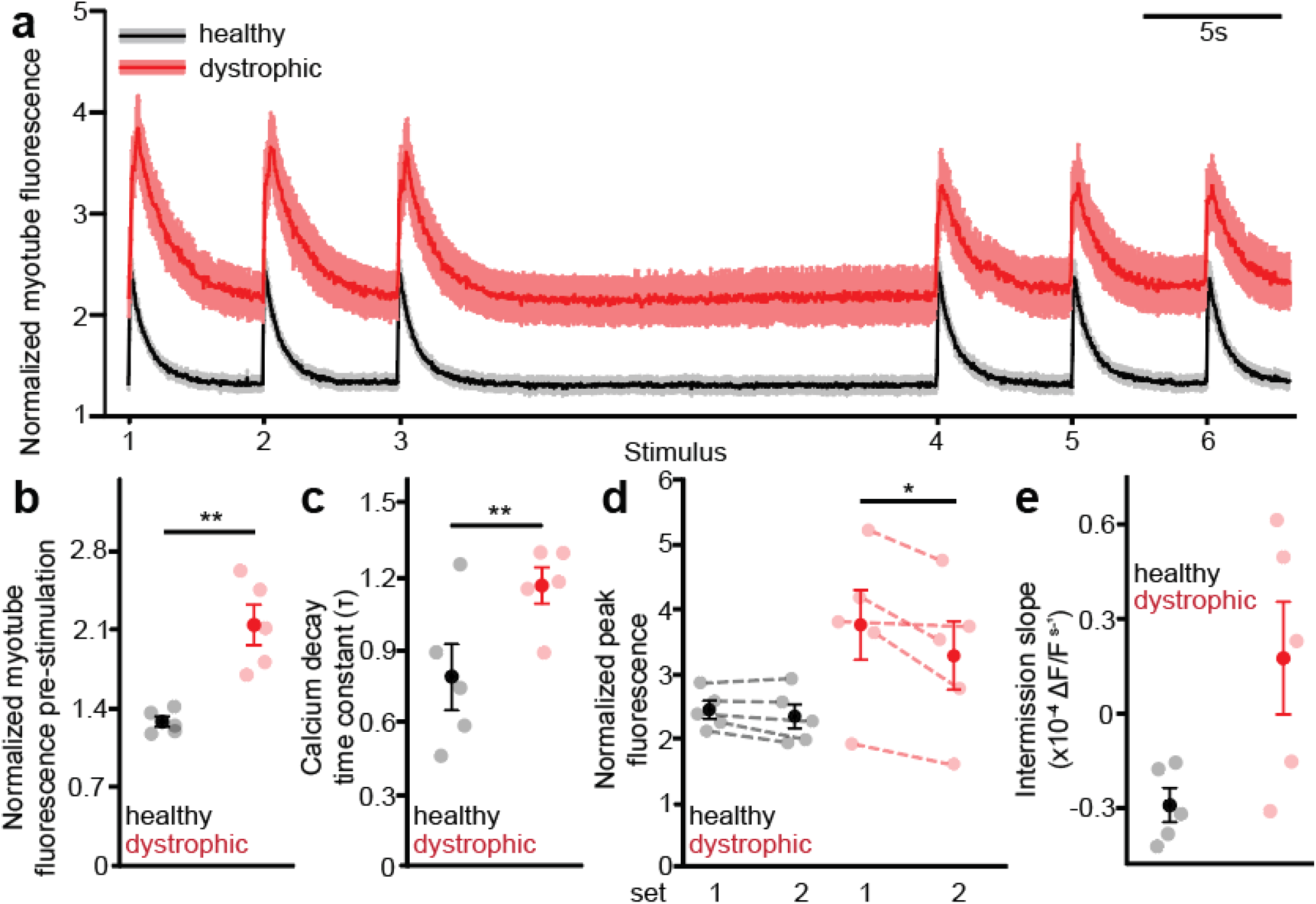
Dystrophic myotubes show elevated baseline calcium and slower calcium clearance. **a)** Average fluorescence traces from healthy (black) and dystrophic (red) myotubes during two sets of repeated electrical stimuli. Traces are of each cell’s own pre-stimulus baseline normalized to the background to highlight dynamics across repeated pulses. **b)** Baseline myotube fluorescence prior to stimulation is higher in dystrophic myotubes (Student’s unpaired t-test). **c)** Calcium decay time constant (τ) following each stimulus (fit with a single exponential) was larger in dystrophic myotubes (Student’s unpaired t-test). **d)** Peak myotube fluorescence during the first and second stimulation sets. The difference in peak amplitude between sets did not reach statistical significance in either genotype (Student’s unpaired t-test). **e)** Intermission slope during the inter-set rest interval, measured in ΔF/F s⁻¹. Positive slope indicates incomplete calcium recovery between sets. The genotype difference trended in the increased direction but did not reach significance (Welch’s t-test, used given unequal variances). Transparent points represent independent replicates, opaque points indicate group means with error bars (mean ± SEM). n = 5 independent replicates of 20 myotubes per group. Significance is indicated (*p < 0.05, **p < 0.01).

### PMCA inhibition during stimulation shifts healthy myotubes dynamics toward dystrophic ones but causes only limited additional worsening in dystrophic myotubes

The myotube data established that dystrophic cells begin with a higher calcium baseline and clear calcium more slowly under repeated stimulation, consistent with the previous finding that PMCA expression and calcium efflux is reduced in dystrophic myotubes. This raised the question of whether reduced PMCA activity alone is sufficient to impose the dystrophic phenotype during repeated activation or whether additional mechanisms are contributing. To test this, we repeated the electrical stimulation protocol in the presence of Caloxin 1B1, reasoning that if impaired PMCA export is the primary driver of the dystrophic calcium dynamics, then acutely inhibiting PMCA in healthy myotubes should shift their response profile toward the dystrophic pattern, while dystrophic myotubes should be less affected because they already operate with constrained calcium export capacity. **Fig. 9a** shows the average responses of healthy and dystrophic myotubes to electrical stimulation in control conditions and in Caloxin 1B1.

**Figure 9.**
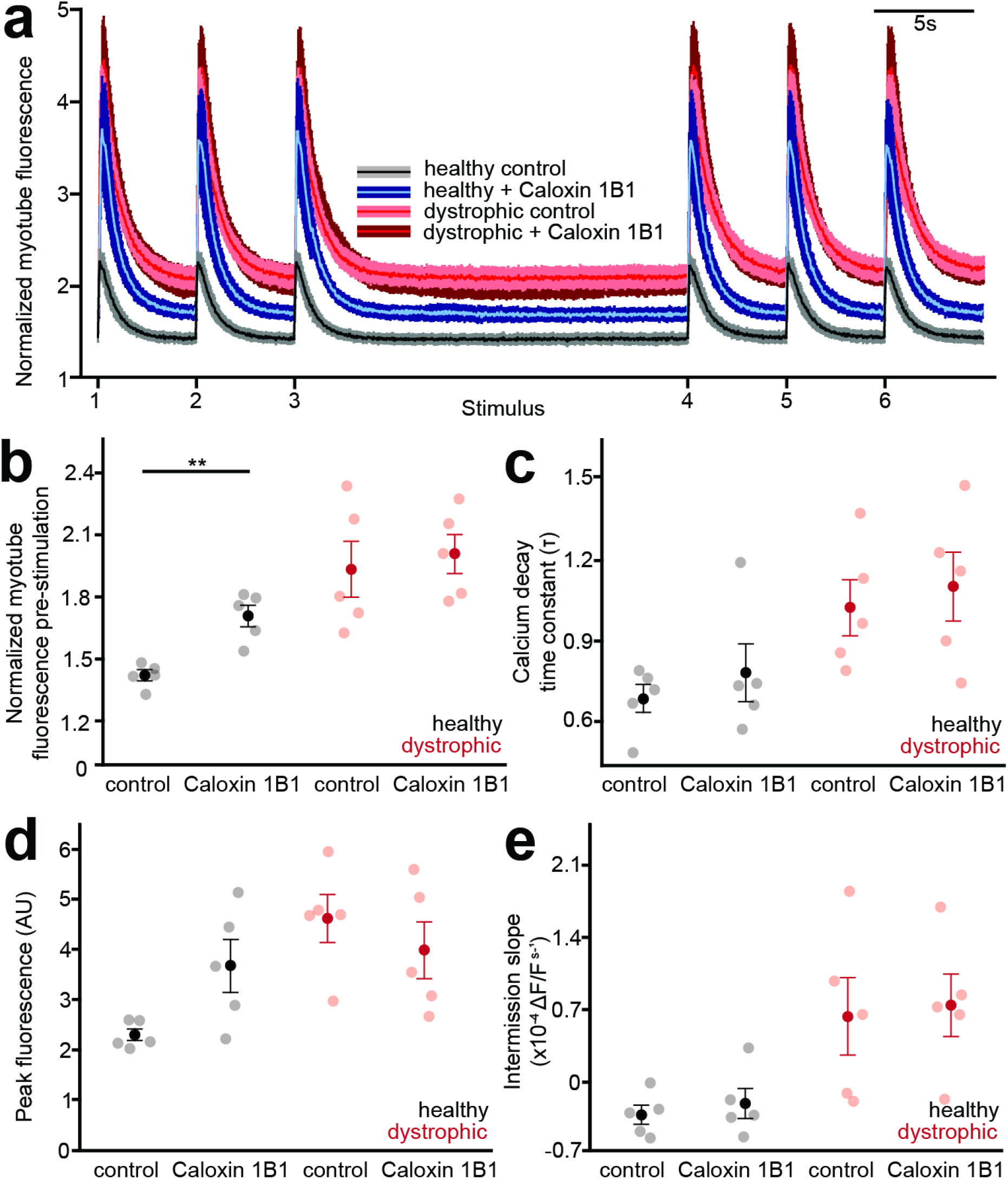
Inhibition of plasma membrane calcium ATPase activity elevates baseline calcium in healthy myotubes toward dystrophic levels, with minimal effects in dystrophic myotubes. **a)** Average normalized fluorescence traces during two sets of repeated electrical stimulation for WT and DMD myotubes treated with vehicle (water) or Caloxin 1B1 (100 µM). **b)** Baseline fluorescence prior to stimulation. Caloxin 1B1 significantly elevated baseline calcium in healthy myotubes, shifting treated healthy myotubes to levels statistically indistinguishable from untreated dystrophic myotubes; the same inhibition did not produce a significant further increase in dystrophic myotubes (Student’s unpaired t-test; Welch’s t-test used for dystrophic comparison given unequal variances). **c)** Calcium clearance kinetics quantified as the efflux time constant (tau) derived from post-stimulus exponential decay fits. Caloxin 1B1 did not significantly slow clearance within either genotype individually, though trends were in the expected direction (Student’s unpaired t-test). **d)** Peak fluorescence responses during the stimulation period. Peak amplitude did not differ significantly within either genotype with Caloxin 1B1 treatment (Welch’s t-test for healthy myotubes, Student’s unpaired t-test for dystrophic myotubes). **e)** Intermission slope during the inter-bout rest interval, measured in ΔF/F s⁻¹. Slope did not differ significantly between vehicle and Caloxin 1B1 within either genotype (Student’s unpaired t-test). The overall pattern across panels supports a model in which reduced PMCA-dependent export capacity in dystrophic myotubes limits the incremental effect of pharmacological inhibition. Transparent points represent independent replicates, opaque points indicate group means with error bars (mean ± SEM). n = 5 independent replicates of 20 myotubes per group. significance is indicated (**p < 0.01).

In healthy myotubes, PMCA inhibition significantly elevated baseline calcium levels (**Fig. 9b**, control: 1.42 ± 0.026; Caloxin 1B1: 1.71 ± 0.052, Student’s t-test, p = 0.001) so that they did not differ significantly from untreated dystrophic myotubes (dystrophic control: 1.93 ± 0.137, Student’s t-test, p = 0.16). This is consistent with PMCA inhibition alone being sufficient to impose a dystrophic-like calcium baseline in healthy cells. In contrast, Caloxin 1B1-induced PMCA inhibition did not affect baseline calcium levels in dystrophic myotubes (dystrophic control: 1.93 ± 0.137, dystrophic + Caloxin 1B1: 2.01 ± 0.095, Welch’s t-test, p = 0.63), consistent with an impaired PMCA-mediated calcium export capacity in dystrophic cells. Caloxin 1B1 did not affect calcium clearance in healthy or dystrophic myotubes (**Fig. 9c**, τ healthy control: 0.686 ± 0.054 s; τ healthy + Caloxin 1B1: 0.780 ± 0.110 s, Student’s t-test, p = 0.46; τ dystrophic control: 0.947 ± 0.113 s; τ dystrophic + Caloxin 1B1: 1.12 ± 0.120 s, Student’s t-test, p = 0.34). Normalized peak calcium fluorescence responses, while exhibiting a nonsignificant but positive trend in healthy myotubes (**Fig. 9d**, healthy control: 2.30 ± 0.122; healthy + Caloxin 1B1: 3.67 ± 0.525, Welch’s t-test, t = -2.54, df = 4.4, p = 0.058) were also unaffected by Caloxin 1B1 treatment (dystrophic control: 4.62 ± 0.476 AU; dystrophic + Caloxin 1B1: 3.98 ± 0.569 AU, Student’s t-test, p = 0.419). Similarly, intermission slope was unaffected by Caloxin 1B1 treatment in both genotypes (**Fig. 9e**, healthy control: −3.19×10⁻⁵ ± 9.40×10⁻⁶ ΔF/F s⁻¹; healthy Caloxin 1B1: −2.08×10⁻⁵ ± 1.46×10⁻⁵ ΔF/F s⁻¹, Student’s t-test, p = 0.544; dystrophic control: 6.35×10⁻⁵ ± 3.77×10⁻⁵; dystrophic Caloxin 1B1: 7.43×10⁻⁵ ± 3.03×10⁻⁵, Student’s t-test, t = −0.223, df = 8, p = 0.829), with no discernible trend in either direction. Together with the absence of a Caloxin 1B1 effect on calcium clearance and peak amplitude, these results suggest that acute PMCA inhibition selectively shifts the resting calcium set point without detectably altering stimulus-evoked calcium dynamics under the conditions used here, indicating that PMCA activity primarily governs basal calcium maintenance rather than recovery kinetics during repeated electrical demand.

### PMCA1 overexpression reduces basal calcium in dystrophic myoblasts with a proportionally greater effect than in healthy cells

The shRNA and pharmacological inhibition experiments established that reducing PMCA activity in healthy cells is sufficient to shift basal calcium levels toward dystrophic levels. However, testing whether a correctly functioning PMCA is also sufficient to restore healthy calcium levels in dystrophic muscles requires the complementary gain-of-function test. We transfected healthy (AB1190) and dystrophic (AB1071) myoblasts with a human codon-optimized PMCA1 overexpression construct fused to mCherry, allowing identification of transfected cells by mCherry fluorescence. We first confirmed that overexpression in transfected cells was successful using antibody staining against PMCA1 in mCherry-positive cells. **Fig. 10a** shows example images for both healthy and dystrophic myoblasts. Both groups showed increased PMCA immunofluorescence in transfected (mCherry-positive) cells relative to untransfected, mCherry-negative controls (**Fig. 10b**, left, healthy control: 36.9 ± 3.32 AU; healthy PMCA1: 91.2 ± 10.2 AU, Student’s t-test, p < 0.001; dystrophic control: 17.6 ± 2.57; dystrophic PMCA1: 52.7 ± 9.35; Student’s t-test, p = 0.007), confirming that the construct increased expression of PMCA1 protein in both genotypes. The proportional PMCA abundance increase was similar between genotypes as reflected by a similar PMCA1-to-control fluorescence ratio (**Fig. 10b**, right, healthy: 2.52 ± 0.327; dystrophic: 2.91 ± 0.261; Student’s t-test, p = 0.379), indicating the overexpression construct drove similar fold changes in PMCA1 despite their different baseline levels.

**Figure 10.**
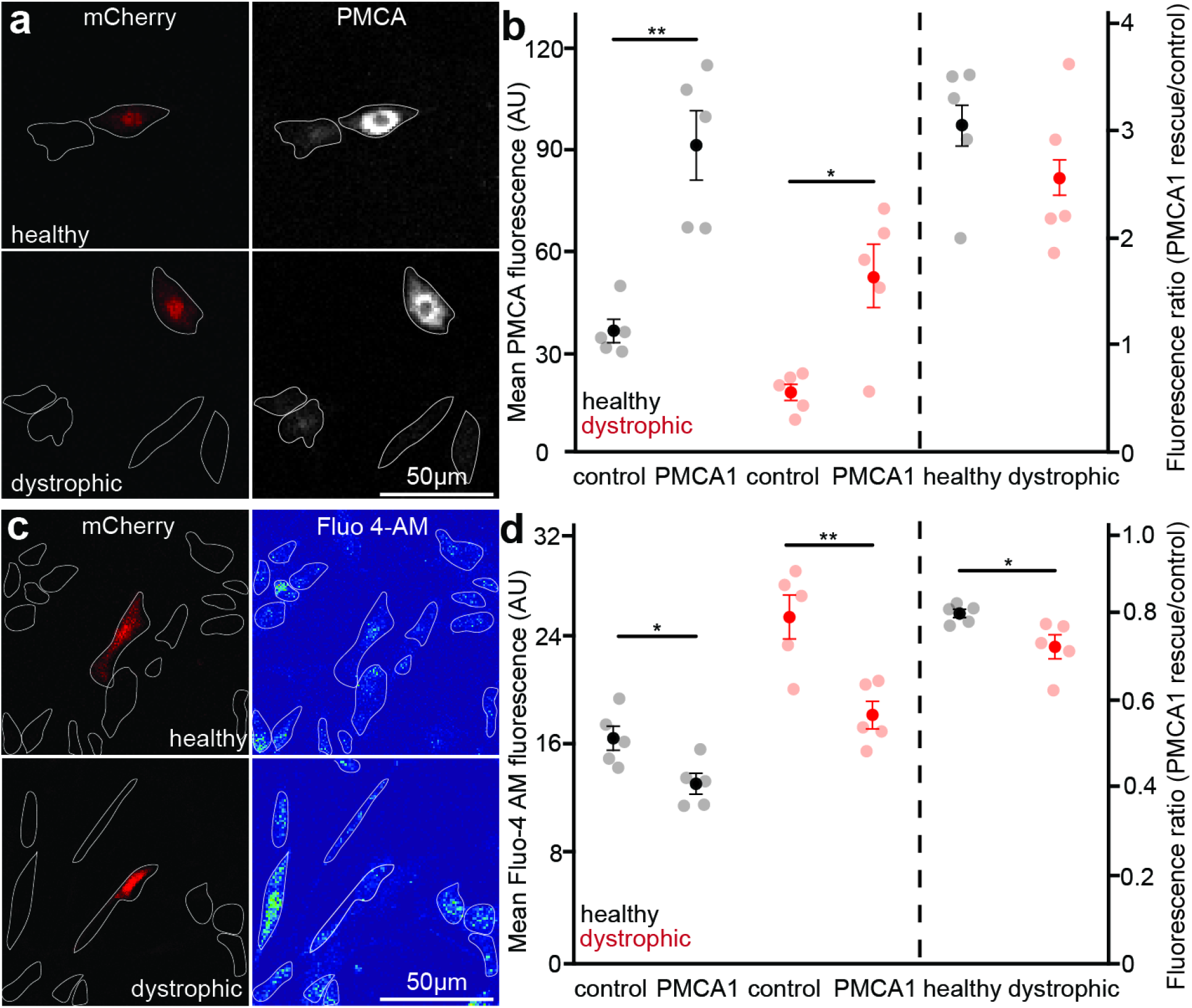
PMCA1 overexpression reduces basal sarcoplasmic calcium in dystrophic myoblasts with a proportionally greater effect than in healthy myoblasts, consistent with restoration of a constrained calcium export reserve. **a)** Representative immunofluorescence images showing mCherry fluorescence (left) and PMCA antibody fluorescence (right) in healthy (AB1190) and dystrophic (AB1071) myoblasts transfected with a PMCA1 overexpression construct, confirming increased PMCA1 abundance in transfected cells. **b)** Quantification of mean PMCA immunofluorescence intensity in mCherry-positive cells from healthy and dystrophic myoblasts. Both genotypes show increased PMCA signal in transfected cells relative to untransfected controls, with dystrophic cells showing a smaller absolute increase consistent with their reduced baseline PMCA abundance (Student’s unpaired t-test). n = 5 independent replicates of 223-712 control cells and 19-54 PMCA1 overexpression cells per group. **c)** Representative images of healthy and dystrophic myoblasts showing mCherry fluorescence (left) to identify transfected cells and Fluo-4 AM signal (right) to visualize basal sarcoplasmic calcium. Transfected cells are identified by mCherry fluorescence and outlined by dashed lines. **d)** Quantification of mean Fluo-4 brightness in control (untransfected) and PMCA1-overexpressing (mCherry+) cells in healthy and dystrophic myoblasts (left axis), and the fluorescence ratio of PMCA1-overexpressing to control cells within each genotype (right axis, separated by dashed line). PMCA1 overexpression significantly reduced basal calcium in both healthy and dystrophic myoblasts, with the proportional reduction being greater in dystrophic cells, consistent with unmet calcium export demand in dystrophic myoblasts (Student’s unpaired t-test). Transparent points represent independent replicates, opaque points indicate group means with error bars (mean ± SEM). n = 5 independent replicates of 174-922 control cells and 13-62 PMCA1 overexpression cells per group. significance is indicated on the plots (*p < 0.05, **p < 0.01).

Having confirmed successful overexpression, we next asked whether the increase in PMCA1 protein was sufficient to reduce basal sarcoplasmic calcium. **Fig. 10c** shows pseudocolor-coded images of healthy and dystrophic myoblasts after Fluo-4 AM application. Basal Fluo-4 AM fluorescence was significantly reduced in PMCA1-overexpressing cells in comparison to untransfected control cells. This was the case in both healthy and dystrophic myoblasts (**Fig. 10d**, left, healthy control: 16.4 ± 0.9 AU; healthy PMCA1: 13.1 ± 0.761 AU; Student’s t-test, p = 0.02; dystrophic control: 25.3 ± 1.61; dystrophic PMCA1: 18.1 ± 1.02; Student’s t-test, p = 0.005). Critically, the proportional reduction in calcium following *PMCA1* overexpression was greater in dystrophic than in healthy myoblasts, as reflected by a smaller PMCA1-to-control fluorescence ratio in dystrophic cells (**Fig. 10d**, right, healthy: 0.797 ± 0.0097; dystrophic: 0.720 ± 0.027; Student’s t-test, p = 0.029). Basal Fluo-4 fluorescence in PMCA1-overexpressing dystrophic cells did not differ from untransfected healthy controls (18.1 ± 1.02 versus 16.4 ± 0.90 AU, difference 1.7 AU, 95% CI −1.4 to +4.8 AU, Student’s unpaired t-test, p = 0.25). Overexpression closed 7.2 of the 8.9 AU genotype difference, about 80 percent, which we report as an interval rather than as equivalence given five replicates per group. This pattern complements the molecular silencing of PMCA and its pharmacological inhibition, together suggesting that dystrophic myoblasts operate with reduced PMCA1 activity and impaired calcium export capacity, which elevates calcium levels in early dystrophic muscle. Rescuing *PMCA1* expression is sufficient to reduce the elevated baseline calcium levels in dystrophic cells toward untreated healthy levels, identifying PMCA1-dependent export as an ongoing determinant of the elevated calcium set point in dystrophic myoblasts.

## Discussion

Although recent work has begun to locate dystrophic abnormalities before birth, calcium dysregulation in Duchenne muscular dystrophy is still overwhelmingly modeled as a mechanical problem, in which loss of dystrophin weakens the link between the cytoskeleton and the extracellular matrix(*36*, *37*). Contractions are thought to strain and tear the sarcolemma, allowing calcium to enter through the resulting lesions. Our data support a two-component account instead: an offset in the resting calcium set point is established in dystrophic muscles before the contractile apparatus is functional, then a second, activity-dependent component is added once contractions begin. This second component amplifies the initial offset rather than creating it. Separating these two components is what *C. elegans* embryos permit, and what mammalian models have not, because it allows calcium to be measured in intact muscles across a developmental window in which the absence of contraction can be independently verified. We found that the early offset is not a fixed consequence of altered development, but an imbalance held in place by a standing deficit in calcium export. Raising export capacity reverses most of it in cells that already are carrying the phenotype. Thus, in our account, Duchenne muscular dystrophy has a developmental origin and a continuous mechanism that moves the earliest addressable point back to the myogenic progenitor (without confining it there).

The early onset component is supported at 400 minutes post-fertilization by three lines of evidence that converge on that timepoint. First, movement is absent by measurement with movement indexes in all groups at or below the 1.2 floor that immobile embryos produce in our system. Second, neuromuscular cholinergic drive is absent by developmental anatomy. Presynaptic vesicle clusters at body wall neuromuscular junctions are not detectable in electron micrographs until approximately 550 mpf, with ventral junctions later still(*38*).

Consequently, at 400 mpf, the machinery for cholinergic drive to muscle has not yet been built, and the contractile apparatus, though assembling, is not yet functional. Third, treatment with d-tubocurarine did not alter the movement index. This confirms the expectation: d-tubocurarine is a nicotinic acetylcholine receptor antagonist, and at a stage with no functional neuromuscular junction, cannot interfere with neurotransmission at said junction. The null effect of dTBC on movement at 400 mpf thus confirms the developmental anatomy rather than indicating a failure of the drug and cannot be interpreted as evidence that movement was already at a floor. Against this background, muscle calcium is nonetheless already elevated in dystrophic embryos, with every dystrophic mean exceeding every healthy one.

This supports the idea that the offset is not a consequence of muscle contraction.

In addition to this early component, a second component appears once movement begins. By 900 minutes post fertilization, dystrophic and healthy embryos move indistinguishably, yet dystrophic embryos showed consistently higher calcium levels. This suggests that similar mechanical input produces unequal calcium accumulation. By this stage the motor program has become synaptically driven, since the mature sinusoidal locomotion pattern emerging after 650 mpf requires synaptic vesicle exocytosis whereas earlier embryonic movement does not(*33*). Blocking cholinergic drive at this timepoint accordingly removes the genotype difference, which is consistent with expectation from an accumulation defect (i.e. removing the input allows steady state to converge due to residual export capacity). This same relationship is evident in the regression of brightness on movement across embryogenesis. There dystrophic embryos show a significantly steeper slope than healthy embryos. A third perturbation behaves differently and is worth setting aside explicitly. Knockdown of the troponin C homolog *pat-10* abolishes movement yet raises muscle calcium in both genotypes. Because troponin C is itself a calcium binding protein, this manipulation removes a cytosolic calcium sink at the same time as it removes contraction, and it therefore cannot be read as a test of the movement hypothesis in either direction. We note it because its direction is opposite to that of the other contractility knockdowns, *lev-11*, *myo-3*, and *unc-38*, all of which reduced both movement and muscle calcium in dystrophic animals, not because it supports the argument. However, we are cautious not to overread the convergence at 900 mpf with three replicates per group, as the 95% confidence interval on that residual difference (treated dystrophic minus untreated healthy, −0.03) spans −0.15 to +0.10 and comfortably contains the offset measured at 400 mpf. Therefore, our data cannot distinguish complete convergence from a persistent baseline difference of the early magnitude.

Re-understanding the phenotype as a displaced setpoint rather than accumulated damage generates a specific mechanistic prediction. Resting cytosolic calcium is produced by the steady state of influx and extrusion. In a pre-contractile cell, the machinery that dominates calcium flux in a mature muscle (principally the ryanodine receptor and SERCA) has not become prominent and the sarcoplasmic reticulum is not yet loaded(*11*, *12*, *14*, *15*). Therefore, plasma membrane handling carries a disproportionate share of the burden of maintaining low resting concentration. Developmental timing alone does not identify which side of that balance is at fault, since resting calcium at any stage reflects influx as well as extrusion, and leak, constitutively active channels and store-operated entry all remain possible contributors. What the timing does is direct attention to basal plasma membrane calcium handling rather than to excitation-contraction coupling. Specifically, our perturbation experiments are consistent with export impairment. Reducing *mca-1* raised muscle calcium in wild type animals towards dystrophic levels but produced no detectable change in dystrophic animals. In contrast, reducing *mca-2* in the same animals under the same model produced a difference, establishing that dystrophic muscle can be driven higher. Thus, the absence of an *mca-1* effect reflects a pathway-specific limitation rather than saturation of the indicator or the cells. A plausible route to such a limitation is suggested by the finding that the dystrophin associated complex organizes distinct planar-polarized membrane compartments at the *C. elegans* sarcolemma, positioning potassium channels and complex components in defined domains established by the end of embryogenesis(*39*). Dystrophin has been reported to modulate the confinement of voltage-dependent calcium channels within sarcolemmal nanodomains in response to varying muscle tonus, which concerns channel dynamics under load rather than resting steady state(*40*). Consistent with that distinction, knockdown of the L type calcium channel *egl-19* did not significantly alter resting muscle calcium in either genotype, which is consistent with entry not being the limiting variable at rest. Our earlier screen provides a direct test of the alternative that reduced sarcoplasmic reticulum uptake, rather than reduced plasma membrane export, sets the dystrophic resting level. Silencing *sca-1* raised basal muscle calcium in healthy and dystrophic animals alike, by a comparable proportion in each (P = 0.009 and P < 0.001, respectively)(*24*), as *mca-2* silencing does here. Only *mca-1* silencing was asymmetric, raising basal calcium in healthy animals by a similar proportion while leaving dystrophic animals unchanged. Were SR uptake the limiting step in dystrophic muscle, further impairing SERCA should have been occluded there; instead the occlusion is confined to *mca-1* among the four clearance pathways tested across the two studies. That the dystrophic level can also be driven back down, as calmodulin knockdown did when it returned basal and peak brightness to healthy values without affecting healthy animals(*24*), is itself consistent with a displaced set point rather than irreversible accumulated damage.

Human myogenic cultures reproduce the pattern in cells that have never contracted and have no assembled sarcomeres. Dystrophic myoblasts showed elevated basal calcium alongside reduced PMCA abundance. Both pharmacological inhibition with Caloxin 1B1, and stable silencing of *ATP2B1* raised basal calcium in healthy cells while producing little further change in dystrophic cells. One caveat is that loss of function experiments of this kind cannot alone distinguish a real deficit in export capacity from a readout in which dystrophic cells simply have less room to respond. We addressed this alternative in our silencing experiments, which raised calcium significantly in dystrophic myoblasts, showing that these cells can be driven higher and that the absence of a pharmacological effect is not a ceiling.

The same experiments set a limit on how far the result can be pushed, since the magnitude of the silencing effect in healthy cells substantially exceeds the dystrophic baseline rather than approaching it, indicating that experimental knockdown removes more PMCA1 activity than the disease state does. Therefore, our interpretation is that PMCA1 appears necessary for maintenance of the resting setpoint and that its reduction is sufficient to shift that setpoint in the dystrophic direction (not that the reduction measured is entirely responsible for the offset). The gain of function experiments are not subject to the same ambiguity. These experiments predicted that cells carrying unmet export demand should show a proportionally larger correction than cells that do not. Indeed, we found that while PMCA1 overexpression lowered basal calcium in both genotypes, the proportional reduction was significantly greater in dystrophic myoblasts, despite an increase in PMCA1 protein that was equivalent across both genotypes. Quantitatively, we found that PMCA1 overexpression closed most of the genotype difference in basal calcium, bringing dystrophic cells to a level not distinguishable from untreated healthy cells, although with five replicates per group that comparison is underpowered and we report it as an interval rather than as equivalence.

Because overexpression also lowers calcium in healthy cells, our interpretation is that dystrophic cells return to the untreated healthy level, rather than being normalized in absolute terms. This interpretation has mechanistic implications, beyond the fact that it does not rest on a null result. The elevated set point in these myoblasts was already established at the time of transfection, and raising export capacity afterwards reversed most of it. The offset is therefore established early but remains correctable after establishment, with ongoing PMCA1-dependent export capacity accounting for a substantial fraction of it. We are careful not to claim more than this. This establishes reversibility, not causation, and partial rescue, not complete normalization. Reversibility by raising export does not by itself establish that a standing export deficit is what holds the offset in place, since increasing export could compensate for a different primary lesion rather than correcting the causal one. What the experiment does establish is that the early dystrophic calcium state is not beyond reach once it has appeared.

A discrepancy between our sequencing and targeted measurements deserves comment. RNA sequencing did not identify ATP2B1 as differentially expressed in dystrophic myoblasts (log2 fold change −0.11, adjusted p = 0.463), while qPCR on independent biological replicates detected a modest but significant reduction (p = 0.018). This discrepancy may reflect the limited power of three RNA-seq replicates per group to resolve small changes after correction for multiple testing, compared with targeted qPCR using five replicates per group. Differences between independently sampled cultures may also contribute. The central interpretation rests on the combined evidence from PMCA immunoreactivity and functional perturbations rather than on transcript abundance alone.

The timing of the human phenotype raises a problem that the established structural understanding does not yet resolve. PMCA interacts with the dystrophin associated protein complex through syntrophin mediated scaffolding(*41*). The natural expectation would be that loss of dystrophin would destabilize PMCA at the membrane. However, in proliferating myoblasts the mature complex has not yet been assembled, yet reduced PMCA1 abundance is already detectable at the transcript level. Therefore, loss of a scaffold that is not yet present cannot be the whole explanation. This argues for a regulatory rather than purely structural route from dystrophic deficiency to constrained calcium export. We identified two possible scenarios: reduced export capacity could be an upstream event that subsequently shapes complex assembly and differentiation, or alternatively dystrophin could control PMCA expression through pathways independent of complex formation. We remain cautious about the candidates our transcriptomic data identified. Reduced *EGR1* and *FOS* expression is consistent with altered activity dependent transcription. However, the documented regulatory link is between EGR1 and ATP2B2 in neural tissues. No comparable link to ATP2B1 in muscle has been established. Whether EGR1 or FOS directly regulate ATP2B1 expression in muscle remains to be tested, and promoter occupancy and reporter assays would be the natural next step to evaluate this possibility. Another consideration is that the shorter dystrophin isoforms, including Dp71, are expressed in proliferating myoblasts(*19*, *42*) and are not removed by the mutation carried by the patient derived line used here. Therefore, the argument that scaffolding cannot explain the deficit applies to the full-length isoform and the mature complex, but the residual short isoform contribution cannot be excluded without isoform-specific manipulations.

The mechanism by which PMCA1-mediated export is reduced in dystrophic myoblasts remains an open question. The modest reduction in PMCA1 transcript detected by qPCR, alongside the more substantial reduction in PMCA protein detected by immunofluorescence, is consistent with post-transcriptional or post-translational contributions acting alongside reduced expression. One candidate mechanism involves calmodulin, the primary activator of PMCA. Calmodulin binds PMCA at its C terminal regulatory domain and activates it, and this interaction is one of the principal routes by which export capacity is upregulated in response to rising cytosolic calcium. Our RNA sequencing data show dysregulation of CALM1 in dystrophic myoblasts, raising the possibility that altered calmodulin availability impairs PMCA activation independently of any change in PMCA protein abundance, such that PMCA1 is limited by its activator rather than by its abundance alone. This is consistent with prior work from our laboratory showing that silencing the calmodulin homolog *cmd-1* in dystrophic *C. elegans* alleviates the dystrophic calcium phenotype, suggesting that calmodulin-dependent calcium signaling is altered in dystrophic muscle across species. Reduced PMCA1 abundance and impaired calmodulin-dependent activation could therefore act together to constrain calcium export in dystrophic myoblasts.

The observation of elevated resting calcium in dystrophic muscle is not itself novel. Elevated resting sarcoplasmic calcium has been reported in dystrophic myotubes across multiple model systems, including mdx mouse and human myotubes(*4*, *43*), and in dystrophic hiPSC-derived skeletal muscle cells at both the myoblast and myotube stage(*44*), alongside abnormal calcium responses during electrical stimulation. Elevated basal calcium is not universally observed, and some studies have found no significant difference in resting calcium between dystrophic and healthy myotubes(*45*). Altered calcium handling has been similarly reported in dystrophic myoblasts, including in mdx mouse, where transcriptomic work has identified downregulation of calcium handling genes(*31*). Our contribution is not the observation itself but the temporal ordering and the identification of which handling step is constrained. These results extend a body of work that has increasingly located dystrophic abnormalities in undifferentiated myogenic cells.

Studies in *mdx* myoblasts have reported broad remodeling of the calcium signaling toolkit, increased metabotropic purinergic responses, and cell autonomous defects in proliferating progenitors. Pluripotent stem cell models have documented impaired cell junction formation, altered stem cell polarity, and elevated calcium during stimulation of dystrophin deficient myotubes. What has been missing is a direct measurement of calcium in a dystrophic animal before movement begins, alongside with a functional test of whether export capacity rather than entry or storage is the constrained variable. Most directly, examination of mdx mouse embryos found the primary myogenic program intact but secondary myogenesis impaired through reduced muscle stem cell polarity, rescued genetically by deletion of AAK1(*36*), and live imaging has revealed impaired migration and precocious symmetric division in dystrophic muscle stem cells(*37*). These studies locate a developmental defect in the progenitor compartment by routes independent of calcium. Our contribution is complementary and consists of two elements in combination: a direct measurement of calcium in a dystrophic animal before contraction is possible, and a functional test of which calcium handling step is constrained. The developmental phenotypes we observe in dystrophic embryos, namely delayed hatching, extended retention of late-stage embryos in utero, and rescue of hatching timing by eggshell weakening, have not, to our knowledge, been described in dystrophin deficient nematodes, where the reported phenotypes are hyperactivity, hypercontraction and exaggerated head bending, with overt degeneration requiring a sensitizing background. They indicate that dystrophin loss carries functional consequences during embryogenesis itself. The retention phenotype, however, has multiple possible origins. Egg laying in this species is governed by a dedicated neuromuscular circuit that our experiments did not isolate from body wall muscle function, and calcium activated potassium channels are a principal determinant of retention in that circuit. Because the dystrophin associated complex is required for correct localization of the BK channel SLO-1 in *C. elegans* muscle(*46*), a circuit level origin for the retention phenotype is a live alternative that our data do not exclude.

Several limitations provide boundaries for our conclusions. First, the human arm of our experiments rests on a single pair of immortalized lines derived from different donors, so genotype and donor background are confounded in any comparison between lines(*47*). The perturbation experiments are internally controlled, in that each line serves as its own reference. Furthermore, the convergence with an evolutionarily distant animal model provides independent support. Attribution of the deficit to dystrophin loss could be further strengthened by inclusion of additional isogenic corrected lines. Secondly, both Fluo-4 and GCaMP2 report relative rather than absolute calcium and are not ratiometric. Therefore, our inferences are restricted to comparisons made within matched acquisition sessions. The specific concern that reporter expression rather than calcium could drive the worm phenotype has been previously addressed for these strains: quantitative PCR found no difference in *myo-3* transcript abundance between genotypes in L1 larvae, although that control was performed at the larval stage and does not by itself extend to the embryonic timepoints reported here. Furthermore, the PMCA antibody available to us does not discriminate between PMCA1 and PMCA4, so a contribution from PMCA4 cannot be excluded. A further limitation concerns the human knockdown experiments. Because the shRNA construct expresses mCherry and the puromycin resistance gene from a single transcript, mCherry-negative cells that survived selection may carry the construct below the detection threshold rather than lack it, so the internal control population may itself be partially silenced. Any such contamination would reduce the measured difference rather than create it. We also did not confirm ATP2B1 knockdown at the transcript or protein level, so the genetic loss-of-function result rests on reporter identity alone and is interpreted alongside, rather than independently of, the pharmacological data. Relatedly, our genetic and pharmacological evidence identifies constrained PMCA-dependent export as a contributor to the elevated set point but does not exclude a concurrent abnormality in calcium influx, which our experiments were not designed to detect. In our dTBC experiments at 900 mpf our three replicates per group were underpowered for the residual comparison we discussed above. Lastly, although dystrophic myotubes clear calcium more slowly under repeated stimulation, which is a functional consequence of the elevated set point, we have not shown that restoring PMCA1 improves muscle function or delays degeneration. The causal link between the shifted set point and degeneration therefore remains inferred rather than demonstrated.

These findings point to several priorities for future work. The most direct test is whether restoring export capacity reduces muscle degeneration, and the *C. elegans* burrowing and exertion assays provide a tractable system in which to test this. It also remains to be determined whether the impaired calcium clearance observed in dystrophic myotubes under repeated stimulation is a direct consequence of the elevated baseline or reflects additional independent defects, since acute PMCA inhibition reproduced the baseline shift but not the kinetic changes. Finally, identifying the upstream regulatory mechanism that reduces *PMCA1* expression in dystrophic myoblasts is likely a more precise therapeutic target than the pump itself, since broadly augmenting PMCA1 activity across cell types would have effects beyond muscle that this study did not examine. Dystrophic muscle carries an elevated calcium set point linked to constrained plasma membrane export from the earliest stages of myogenesis in two species; that offset precedes the mechanical events usually invoked to explain it; mechanical activity thereafter amplifies rather than creates it; and it is sustained by an ongoing export deficit rather than fixed during development, so that restoring export capacity substantially reverses it. Calcium dysregulation in Duchenne muscular dystrophy is on this account developmental in origin and continuous in mechanism: the earliest addressable point lies earlier than the current framing implies, and the deficit persists rather than resolving as the muscle matures.

## Methods

### *C. elegans* strains and culture conditions

Animals were maintained at 20°C on nematode growth media (NGM) agar plates seeded with *Escherichia coli* OP50 as previously described(*48*). The wild-type N2 and dystrophic BZ33 (*dys-1(eg33*)) strains were obtained from the Caenorhabditis Genetics Center (CGC; supported by NIH Office of Research Infrastructure Programs, P40 OD010440). The calcium-reporter strain ZW495 (*zwIs132[myo-3p::GCaMP2 + lin-15(+)])* was also obtained from the CGC. The dystrophic reporter strain AVG6 (*dys-1(eg33*); *zwIs132[myo-3p::GCaMP2 + lin-15(+)])* was generated by crossing ZW495 with BZ33 and confirmed by PCR genotyping. Strains used in swimming and burrowing assays were maintained on plates supplemented with concentrated OP50 to ensure adequate nutrition during behavioral testing.

### Egg synchronization and embryonic staging

To obtain developmentally staged embryos, day-one adult hermaphrodites were allowed to lay eggs for one hour at 20°C on seeded NGM plates. Adults were then removed, and only eggs at or prior to the two-cell stage were retained for analysis. Eggs more advanced than expected for the 150-minute post-fertilization stage were excluded prior to imaging to ensure a uniform developmental window.

### Hatching curve analysis

Hatching kinetics were quantified by fitting a three-parameter sigmoid function to cumulative hatching data for each experimental group using SigmaPlot 11.0 (Systat Software). The equation used was f = a/(1 + exp(−(x − x0)/b)), where a represents the upper asymptote (maximum fraction hatched), x0 represents the inflection point (time post-fertilization at which 50% of embryos had hatched, in minutes), and b represents the hatching spread (a time constant in minutes describing the duration of the hatching window, where larger values indicate more dispersed hatching kinetics). Parameters were estimated using dynamic fitting with 200 iterations and 200 fits per group. The T50 (x0, defined as the timepoint in which half of all embryos have hatched) and hatching window (b, defined as the duration of the hatching window over which the majority of embryos transition from unhatched to hatched) were extracted from the best-fit solution for each replicate and compared between groups using Student’s unpaired t-test as described in the Statistical analysis section.

### Embryonic movement quantification

Embryonic movement was quantified from time-lapse recordings acquired at 30 frames per second using a custom pixel-variance approach in ImageJ(*49*). For each embryo, the longitudinal axis was defined as the major axis of the ellipsoidal eggshell, oriented from the anterior to posterior pole. Pixel intensity variance was calculated along this axis over a five-second window (∼150 frames). The resulting variance was normalized to background variance measured immediately outside the eggshell in a matched region, yielding a dimensionless movement index that reflects relative embryo activity independent of posture. Because the brightness ratio is calculated as embryo pixel variance divided by background pixel variance, a value of 1.0 would indicate that the embryo has identical variance to the background, reflecting the theoretical minimum of no movement. In practice, immobile embryos yield a ratio slightly above 1.0, typically around 1.2, due to residual optical noise, minor focal drift, and low-level fluorescence heterogeneity within the eggshell that exceed background variance even in the absence of detectable movement. This baseline ratio serves as the empirical minimum for non-moving embryos in our system. True embryonic movement onset, established by direct visual observation of the first detectable twitch, occurs at approximately 430 minutes post-fertilization. Because early, low-amplitude twitches often fall within the empirical noise floor of the pixel-variance index (∼1.2), the index does not reliably register movement above background until approximately 480 minutes post-fertilization; the 480 mpf value used elsewhere in this manuscript therefore reflects the detection threshold of our quantitative movement metric, not the true onset of muscle activity No spatial or temporal smoothing or intensity thresholding was applied prior to variance calculation.

### Calcium imaging in *C. elegans* embryos

Calcium imaging was performed in animals carrying the muscle-specific reporter *zwIs132[Pmyo-3::GCaMP2::unc-54 3’UTR]* in either wild-type (N2) or *dys-1(eg33)* backgrounds, as previously described(*24*, *50*). Motion recording was followed by single-frame fluorescence imaging under blue-light illumination using a Sola Light Engine at 90x magnification and 30ms exposure time. Images were acquired to minimize fluorophore bleaching. Identical illumination and acquisition settings were used across genotypes and treatments.

### Calcium quantification in *C. elegans*

Fluorescence quantification was performed in ImageJ. A region of interest (ROI) was drawn around each egg or animal, and the maximum GCaMP2 pixel intensity within the ROI was recorded as the activity metric. Peak brightness was used in preference to mean intensity because it reliably tracks the brightest muscle band within each frame and is less sensitive to posture-dependent variation. A matched ROI of equal area positioned immediately adjacent to the egg or animal served as background. The brightness ratio was calculated as egg (or animal) fluorescence divided by background fluorescence, providing a normalized measure that controls for illumination variance between imaging sessions.

### *C. elegans* RNA interference

RNAi-mediated gene knockdown was performed by bacterial feeding. Wild-type (ZW495) and dystrophic (AVG6) animals were raised on NGM plates containing 50 µg/mL ampicillin and 1 mM IPTG, seeded with HT115(DE3) bacteria expressing either the L4440 empty vector (negative control) or dsRNA targeting genes of interest (*mca-1, mca-2, lev-11, myo-3, pat-10, unc-38, egl-19, unc-2,* or *chs-1*). Gravid adults were transferred to RNAi plates and their progeny were used for all analyses. Animals were scored after one day of RNAi exposure (L1 larval stage) or three days (day-one adults). Knockdown efficiency was confirmed by quantitative RT-PCR for *mca-1* and *chs-1*; the remaining targets (*lev-11, myo-3, pat-10, unc-38, egl-19, unc-2*) were validated by the phenotypic outcomes observed (reduced swimming frequency and muscle fluorescence consistent with prior reports).

### *C. elegans* swimming assays

Swimming frequency was measured in day-one adults. Individual animals were placed within a 1 cm copper ring on an NGM agar plate and flooded with 250 µL liquid NGM. Animals were allowed to acclimate for 30 seconds before recording. Locomotion was filmed for 10 seconds and swimming frequency was calculated as the number of complete body bends per second over the recording window.

### Egg paralysis assays

To pharmacologically suppress embryonic movement without inducing excitatory calcium entry, d-tubocurarine chloride (dTBC, Sigma, cat. #5051450001), a competitive antagonist of nicotinic acetylcholine receptors, was administered maternally. Day-one gravid hermaphrodites, which predominantly carry early-stage embryos, were placed on NGM plates seeded with OP50 and supplemented with 1 mM dTBC. Exposure continued until full adult paralysis was confirmed by visual inspection, indicating sufficient internal drug concentration. Paralyzed adults were bleached within 30 minutes of visible paralysis using an alkaline hypochlorite solution to dissolve maternal tissue and release embryos. This timing minimized drug exposure beyond the paralysis endpoint while preventing embryo development past the target stage. Following bleaching, embryos were rinsed thoroughly in NGM buffer and transferred to unseeded NGM agar plates. No dTBC was added to these imaging plates. Suppression of embryonic movement during subsequent incubation and imaging relied on drug retained in the embryos following animal exposure.

Embryos were incubated at 20°C for 150 minutes, corresponding to the time required for fertilized embryos at the time of maternal paralysis to reach the late gastrula stage at which they would have been laid under normal conditions. After incubation, embryos at the expected gastrula stage were visually identified and selected for downstream imaging and analysis. Only embryos at the gastrula stage were included to ensure consistency in developmental timing and dTBC exposure duration. Suppression of embryonic movement by dTBC was confirmed quantitatively using the pixel-variance motion metric described above.

### Cell Culture

Immortalized healthy AB1190 and dystrophic AB1071 myoblasts were kindly provided from the Center for Research in Myology, Sorbonne Université-INSERM, Paris, France. Cells were maintained in standard growth conditions in Skeletal Muscle Growth Medium (PromoCell, cat. #C-23060) and grown on 1:100 Matrigel-coated plasticware (Corning, ref. #356231, lot #3060004) coated plates. For experiments, 2×10^5 cells/cm^2^ were seeded and upon reaching confluency ∼2 days later, growth medium was replaced with a differentiation medium consisting of DMEM (Sigma, cat. #D6046), 1% insulin-transferrin-selenium (Thermo, cat. #41400045) and 2% human serum (Sigma, cat. #H6914). Myoblasts were allowed to differentiate for 48 hours prior to use in experiments.

### Ethics statement

The human myoblast lines used in this study were derived by the MyoLine platform from muscle biopsies obtained with donor informed consent under the ethical approvals held by the Institut de Myologie.

### Human myoblast calcium imaging

Basal cytosolic calcium was measured in undifferentiated myoblasts approximately 8 hours after seeding, prior to the onset of differentiation. Cells were loaded with 5 µM Fluo-4 AM (Thermo Fisher, cat. #F14201) in growth medium for 30 minutes at 37°C, then washed and replaced with fresh live imaging media for microscopy. Confocal imaging was performed using a Leica SP8 laser scanning confocal microscope equipped with an Acousto-Optical Beam Splitter (AOBS). Images were acquired using a 10x/0.4 NA objective. Fluo-4 was excited with a white light laser at 488 nm, and emission was collected on a PMT from 500 to 545 nm. The pinhole was set to 1.0 Airy Unit for all channels. Images were acquired at 512 × 512 pixels (pixel size 2.27 µm; field of view 1,162.5 × 1,162.5 µm) with a scan speed of 400 Hz and a line average of 4. Individual myoblasts were identified by morphology and manually segmented using ROIs drawn in ImageJ. Mean Fluo-4 AM fluorescence intensity was recorded per cell, measured in Arbitrary units (AU). Between 150 and 200 cells were analyzed per biological replicate, sampled from at least 3 fields of view per well. Five independent biological replicates were performed for each condition, each representing a separate plating from independently maintained cultures.

### Human myotube calcium imaging and electrical stimulation

For differentiated myotube experiments, cells were loaded with 5 µM Fluo-4 AM in growth medium for 30 minutes at 37°C, washed twice with phenol red-free, HEPES-buffered live imaging medium (Thermo Fisher, cat. # A59688DJ) and imaged immediately in the same buffer on a Leica SP8 confocal microscope (as described above). Individual myotubes were identified by morphology (multinucleated, elongated cells) and manually segmented using ROIs drawn in ImageJ. Between 19 and 20 myotubes were analyzed per biological replicate across five independent experiments per condition. Calcium dynamics were assessed during two bouts of field electrical stimulation delivered via platinum electrodes placed in the imaging chamber at 15 V. Each stimulation bout consisted of five pulse trains, each 200 ms in duration at 100 Hz, delivered once every 5 seconds over a total bout duration of 20 seconds. The two bouts were separated by an approximately 20-second intermission period, which was timed manually. Fluorescence was recorded for 5 seconds prior to the first stimulus to establish a pre-stimulus baseline. Fluorescence traces were quantified by each cell’s own pre-stimulus baseline normalized to the background, defined as the maximum fluorescence intensity immediately preceding the first stimulus to facilitate comparison of calcium dynamics across cells and conditions. Calcium clearance kinetics were quantified by fitting a single exponential decay function to the post-stimulus fluorescence decline for each pulse train, and the time constant tau was extracted as the primary clearance metric, where a larger tau reflects slower calcium removal. Peak fluorescence was defined as the maximum intensity reached during each stimulation bout. Intermission slope was calculated as the linear regression slope of fluorescence during the inter-bout rest interval, where a positive slope indicates incomplete calcium recovery between bouts. For pharmacological inhibition experiments, Caloxin 1B1 (MedChemExpress, cat. #HY-P5421) was dissolved in water and applied at a final concentration of 100 µM in live imaging medium for 30 minutes prior to the start of electrical stimulation. Water-treated cells of both genotypes served as vehicle controls. Caloxin 1B1 was present throughout both stimulation bouts and the intervening intermission.

### Human myocyte RNA interference

Stable PMCA1 (ATP2B1) knockdown was achieved using the PiggyBac transposon system (VectorBuilder). A construct expressing shRNA targeting ATP2B1 (pPB[shRNA]-mCherry:T2A:Puro-U6>hATP2B1[shRNA#1]) was co-transfected with a transposase expression plasmid (pRP[Exp]-EGFP/Puro-CAG>PBase) into myoblasts using Lipofectamine 3000 (Thermo Fisher) according to the manufacturer’s instructions. Transfected cells were selected with 5 µg/mL puromycin in growth medium for 3 days, after which puromycin was withdrawn and cells were maintained in standard growth medium for an additional 4 days before use in experiments, for a total of 7 days post-transfection. This interval allowed sufficient time for stable transgene expression, PMCA1 protein turnover, and degradation of the transiently expressed EGFP-tagged transposase plasmid; the latter was necessary to prevent spectral bleed-through into the Fluo-4 detection channel during subsequent calcium imaging. PMCA1 shRNA-expressing cells were identified by mCherry fluorescence. mCherry-negative cells in the same cultures, which underwent identical transfection, puromycin selection, Fluo-4 loading and imaging, served as internal controls. Only cells that remained attached and spread with normal myoblast morphology at day 7 were analyzed; rounded or detached cells were excluded. mCherry-positive cells were less numerous than mCherry-negative cells in the same dishes (17-71 versus 175-391 per replicate in healthy cultures, and 8-56 versus 114-544 in dystrophic cultures). Control cells were therefore puromycin survivors without detectable mCherry rather than naive untransfected cells. All imaging was performed on a Leica SP8 confocal microscope under identical acquisition settings across conditions as described above mCherry was excited with a white light laser at 561 nm, and emission was collected from a PMT from 575 to 635 nm.

### Human myocyte PMCA1 overexpression

PMCA1 (ATP2B1) overexpression was achieved using a construct expressing human codon-optimized ATP2B1 (NM_001366525.1) fused to mCherry under CMV promoter control (pRP[Exp]-mCherry-CMV>{hATP2B1[NM_001366525.1](hu co)}, VectorBuilder cat. # VB260324-1121zpp). Transfection was performed via Lipofectamine 3000 (Thermo Fisher, cat. #L3000001) according to the manufacturer’s instructions and cells were allowed to grow for 48 hours before imaging. Transfected cells were identified by mCherry fluorescence and used for downstream calcium imaging and PMCA immunofluorescence as described above.

Overexpression was confirmed at the protein level by PMCA immunofluorescence in mCherry-positive cells relative to untransfected controls. All imaging was performed on a Leica SP8 confocal microscope under identical acquisition settings across conditions as described above

### Immunofluorescence microscopy

Human myoblasts cultured on Matrigel-coated 18 mm glass coverslips were fixed in 10% neutral-buffered formalin for 15 minutes at room temperature, permeabilized with 0.1% Triton X-100 in PBS for 5 minutes at room temperature, and blocked with 0.2% fish skin gelatin in PBS for 5 minutes at room temperature. Primary antibody against PMCA1 and PMCA4 (Thermo Fisher, cat. #PA5-107073; 1:50 dilution in blocking buffer) was applied for 1 hour at room temperature.

Coverslips were then washed three times with PBS for 5 minutes per wash before application of secondary antibody (Alexa Fluor 647-conjugated anti-rabbit IgG, Thermo Fisher, cat. #A-A32733TR; 1:500 dilution in blocking buffer) for 1 hour at room temperature. Nuclei were counterstained with DAPI, and coverslips were mounted with ProLong Gold antifade reagent (Thermo Fisher, cat. #P36930) and allowed to cure overnight at room temperature before imaging.

All imaging was performed on a Leica SP8 confocal microscope under identical acquisition settings across conditions as described above. Alexa Fluor 647 was excited with a white light laser at 633 nm, and emission was collected from a PMT from 650 to 720 nm. Mean PMCA immunofluorescence intensity was quantified by morphology and manually segmented using ROIs drawn in ImageJ. Mean fluorescence intensity was recorded per cell, measured in Arbitrary units (AU). Five independent biological replicates were performed per condition, each representing a separate plating from independently maintained cultures.

### RNA extraction and cDNA synthesis

Total RNA was extracted from five independent biological replicates per condition for both *C. elegans* and human myoblast samples. For *C. elegans*, day-one adults were collected into TRIzol reagent (Thermo Fisher, cat. #15596026) and immediately snap-frozen in liquid nitrogen. RNA was then extracted following the manufacturer’s protocol. For human myocytes, total RNA was extracted from cell cultures using the Direct-zol RNA Miniprep Kit (Zymo, cat. #R2051) according to the manufacturer’s instructions. RNA quality and concentration were assessed for all samples by NanoDrop 2000 spectrophotometry prior to downstream use. For each biological replicate, 500 ng of total RNA was reverse transcribed in an independent cDNA synthesis reaction using the RT Premix Kit (Zymo, cat. #R3012) according to the manufacturer’s protocol.

### Quantitative PCR

Quantitative PCR (qPCR) was performed on each independently synthesized cDNA using PowerUp SYBR Green Master Mix (Thermo Fisher, cat. #A25742) on a QuantStudio 7 Real-Time PCR System (Applied Biosystems). Each 10 µL reaction contained 5 µL PowerUp SYBR Green Master Mix, 0.5 µM each of forward and reverse primers (sequences listed in Table S1), and 1 µL cDNA template. Thermal cycling conditions were as follows: UDG activation at 50°C for 2 minutes, polymerase activation at 95°C for 2 minutes, followed by 40 cycles of denaturation at 95°C for 15 seconds and annealing/extension at 60°C for 1 minute. A melt-curve analysis was performed after amplification (60°C to 95°C at 0.1°C/s with continuous fluorescence acquisition) to confirm amplicon specificity. Each sample was run in technical triplicate, and mean Ct values across triplicates were used for analysis. Relative gene expression was calculated using the ΔΔCt method. Expression was normalized to the housekeeping gene *pmp-3* in *C. elegans* and *RPL13* in human myoblasts. For knockdown validation experiments, expression in RNAi-treated animals was normalized to the L4440 empty-vector condition within each replicate. Data were analyzed using QuantStudio Design and Analysis Software (Applied Biosystems), and statistical comparisons were performed as described in the Statistical analysis section below.

### RNA sequencing

Total RNA was prepared from three independent biological replicates of healthy (AB1190) and dystrophic (AB1071) myoblast populations. Library preparation, including poly-A selection, RNA fragmentation, and cDNA synthesis, and sequencing were performed by Azenta Life Sciences (Illumina platform, 150 bp paired-end reads, approximately 50 million reads per sample). Raw reads were trimmed for adapter sequences and assessed for quality prior to alignment. Reads were aligned to the human reference genome (GRCh38) using STAR (v2.5.2b). Gene-level read counts were generated using feature counts. Genes with fewer than 10 raw counts across all samples were excluded prior to differential expression testing. Read count distributions were examined before and after normalization. Raw counts were normalized using the median-of-ratios method implemented in DESeq2 to account for differences in sequencing depth between samples. Sample similarity was assessed by Euclidean distance-based hierarchical clustering and principal component analysis of normalized count matrices to identify any outlier samples prior to differential expression analysis. Differential expression analysis was performed in R using DESeq2 with a simple two-group design (healthy versus dystrophic) and no additional covariates. Genes were considered differentially expressed at a Benjamini–Hochberg adjusted p-value < 0.05. Reactome pathway enrichment analysis was performed on the differentially expressed gene set using the full detected gene list as the background universe. Cross-species ortholog mapping for the *C. elegans* comparison in Fig. S1B was performed using WormBase, with overlap defined as the fraction of genes within each human Reactome pathway for which an orthologous worm gene was also differentially expressed in dystrophic *C. elegans*, irrespective of direction of change. Raw sequencing data and processed count matrices are publicly available through the NCBI Gene Expression Omnibus (GEO accession: GSE299627).

### Statistical analysis

All statistical analyses were performed in SigmaPlot 11.0 and/or GraphPad Prism. Data distributions were assessed using the Shapiro-Wilk test together with visual inspection where appropriate. For two-group comparisons of normally distributed data with approximately equal variances, Student’s unpaired t-test was used. Where variance heterogeneity was indicated by visual inspection or formally confirmed by the Brown-Forsythe test, Welch’s t-test was applied instead to account for variance heterogeneity without assuming equal group variances. Non-parametric two-group comparisons were made using the Mann-Whitney U test. For multi-group comparisons, one-way ANOVA followed by Holm–Šídák-adjusted post hoc comparisons was used for normally distributed data, and the Kruskal-Wallis test followed by Dunn’s post hoc comparisons was used for non-normally distributed data. Unless a figure legend states otherwise, the experimental unit (n) is the independent biological replicate, and individual embryos, worms, myoblasts, and myotubes within a replicate were treated as subsamples summarized at the replicate level before testing. Where a legend specifies the individual animal as the test unit (N), that departure is stated in the legend. Unless otherwise indicated, five independent biological replicates were analyzed per condition. Statistical significance was defined as p < 0.05. All data are presented as mean ± SEM unless otherwise noted. The relationship between embryo movement index and normalized fluorescence intensity was analyzed using a single linear regression model including genotype, movement index, and a genotype × movement interaction was fitted via ANCOVA. The interaction was used to test whether the regression slopes differed between genotypes, while the genotype main effect tested differences in regression intercepts. Statistical significance was defined as *p* < 0.05. For qPCR experiments, statistical comparisons were performed on ΔCt values using Student’s paired t-test. Healthy and dystrophic samples were paired by batch: within each replicate the two genotypes were grown on plates side by side, and RNA extraction and cDNA synthesis were carried out together in the same session. Separate replicates were collected on different days from different culture batches. Pairing is therefore by batch and does not imply donor matching, since the two lines derive from different donors. For RNA sequencing data, differential expression significance was assessed using the Wald test within the DESeq2 framework, with p-values adjusted for multiple comparisons using the Benjamini–Hochberg procedure, as described in the RNA sequencing section above. These analyses were performed independently of the behavioral and imaging statistics described here.

## Data availability

RNA sequencing data generated in this study, including raw reads and processed count matrices, have been deposited in the NCBI Gene Expression Omnibus under accession GSE299627. Values underlying all quantified panels in Figs. 1 to 5 and 7 to 10, together with the statistical tests applied, exact p values, and n for each comparison, are provided in Data file S1. Unprocessed measurements from which these values were derived, including per-animal intensity and background values, per-cell fluorescence intensities, qPCR Ct values, and frame-level myotube fluorescence traces, are provided in Data file S2. The pathway enrichment and ortholog overlap table underlying Fig. S1 is provided in the Supplementary Materials. *C. elegans* strains N2, BZ33, and ZW495 are available from the Caenorhabditis Genetics Center; strain AVG6 available from the authors on request. Immortalized human myoblast lines are available from MyoLine platform of the Institut de Myologie (Paris, France), subject to a material transfer agreement. All other data supporting the findings of this study are available within the paper and its Supplementary Materials.

## Code availability

Code used to generate Fig. S1 from the pathway enrichment and ortholog overlap table is provided in the Supplementary Materials. All other analyses used published software cited in Methods.

## Materials availability

The immortalized human myoblast lines AB1190 and AB1071 were obtained from the MyoLine platform, Institut de Myologie, Paris, under a collaboration letter and material transfer agreement, and cannot be redistributed by the authors. Requests for these lines should be directed to the MyoLine platform. *C. elegans* strains generated in this study are available from the corresponding author on request.

## Author contributions

D.M. conceived and designed experiments, performed *C. elegans* and human cell culture experiments, analyzed data, and wrote the original draft. K.H.-W. contributed to *C. elegans* calcium imaging and analysis. W.S. contributed to experimental design, embryonic imaging analysis, and manuscript revision. X.D. designed and performed the statistical analyses. P.J. performed experiments and analyzed data. A.S.-C. provided clinical expertise on Duchenne muscular dystrophy, contributed to experimental design, and contributed to writing and revision of the manuscript. J.O. generated and provided the immortalized human myoblast lines AB1190 and AB1071 and reviewed the manuscript. M.F.E. contributed to the design of the human cell experiments, and to manuscript revision. A.G.V.-G. supervised the project, acquired funding, designed experiments, and wrote and revised the manuscript. All authors read and approved the final manuscript.

## Supplementary

**Fig. S1.**
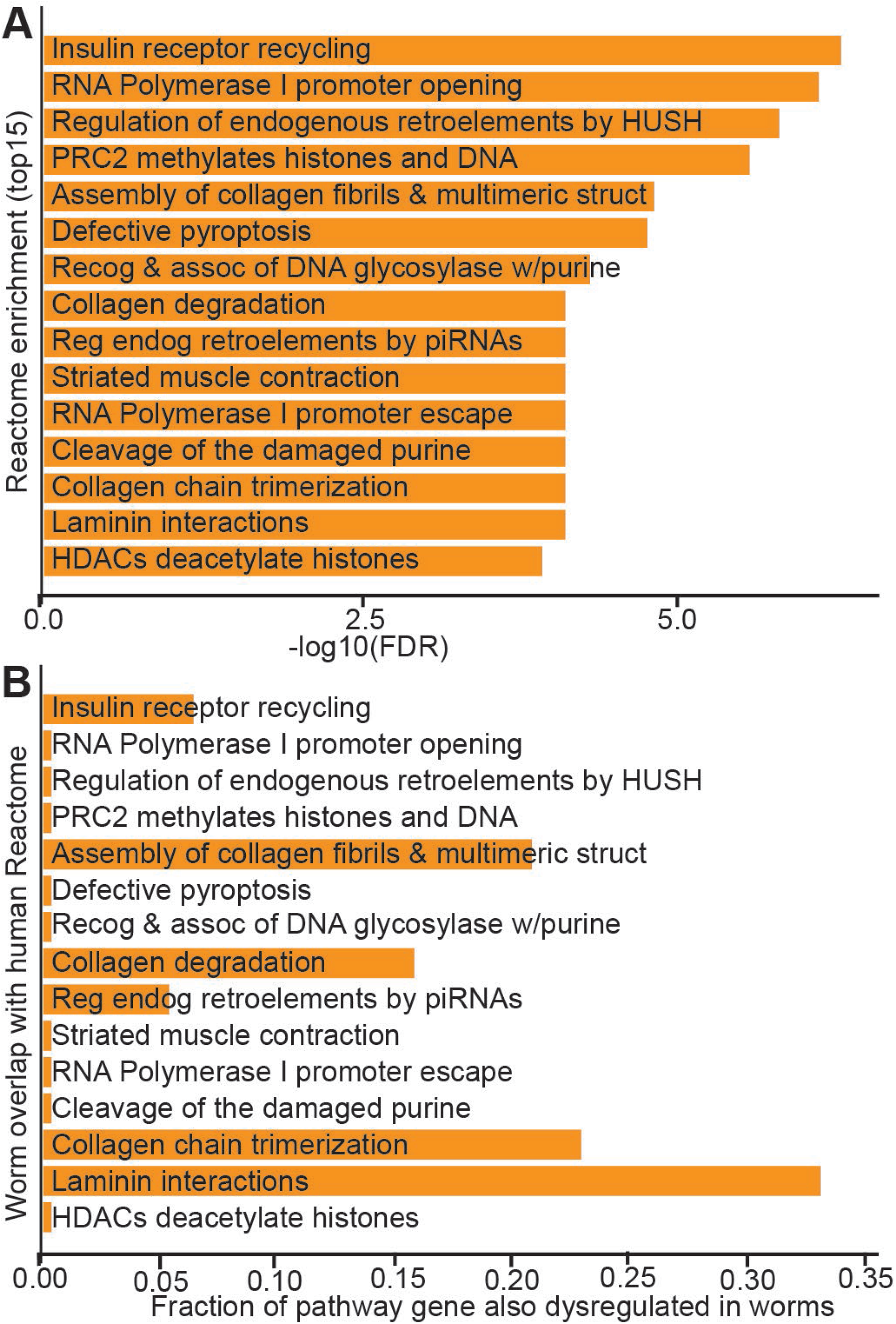
Reactome pathway enrichment in dystrophic human myoblasts and conserved pathway involvement in dystrophic worms. **a)** Top fifteen Reactome pathways enriched among differentially expressed genes in dystrophic versus healthy human myoblasts, ranked by significance and plotted as −log10 of the false discovery rate (FDR). **b)** For the same pathways in the same rank order, fractional overlap with the dystrophic worm transcriptomic signature. Overlap was calculated as the fraction of genes within each human pathway for which an orthologous worm gene was also differentially expressed in dystrophic *C. elegans*, irrespective of direction of change (ortholog mapping via WormBase). Pathways with near-zero values indicate little to no overlap between the human-enriched pathway and the worm differentially expressed gene set.

**Table S1.**
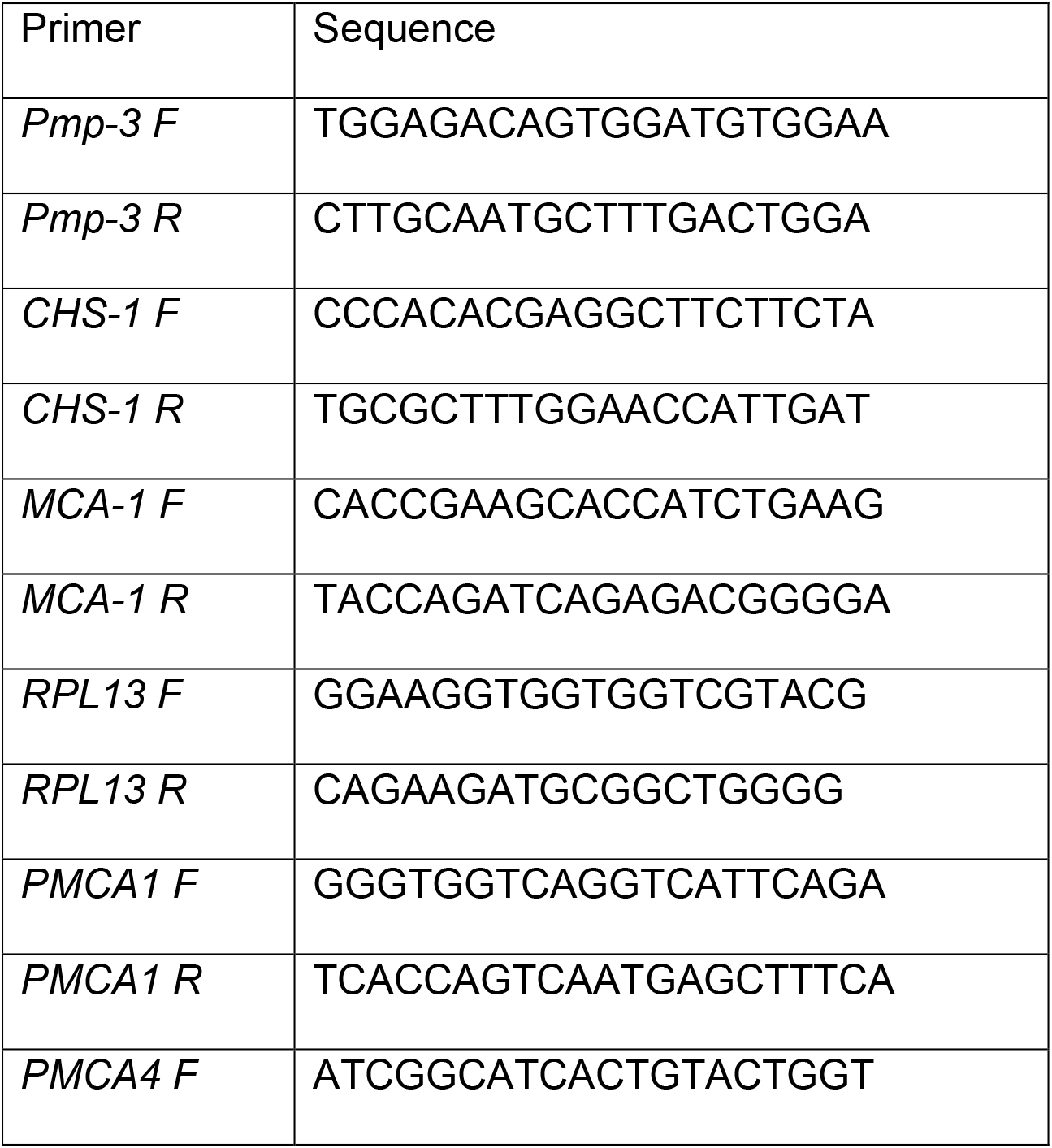
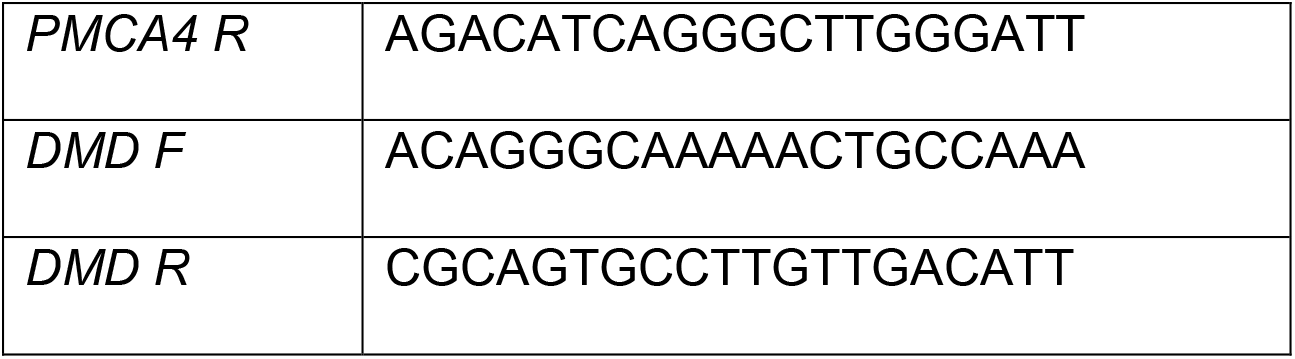
qPCR primer pairs used in this work.

**Table S2.**
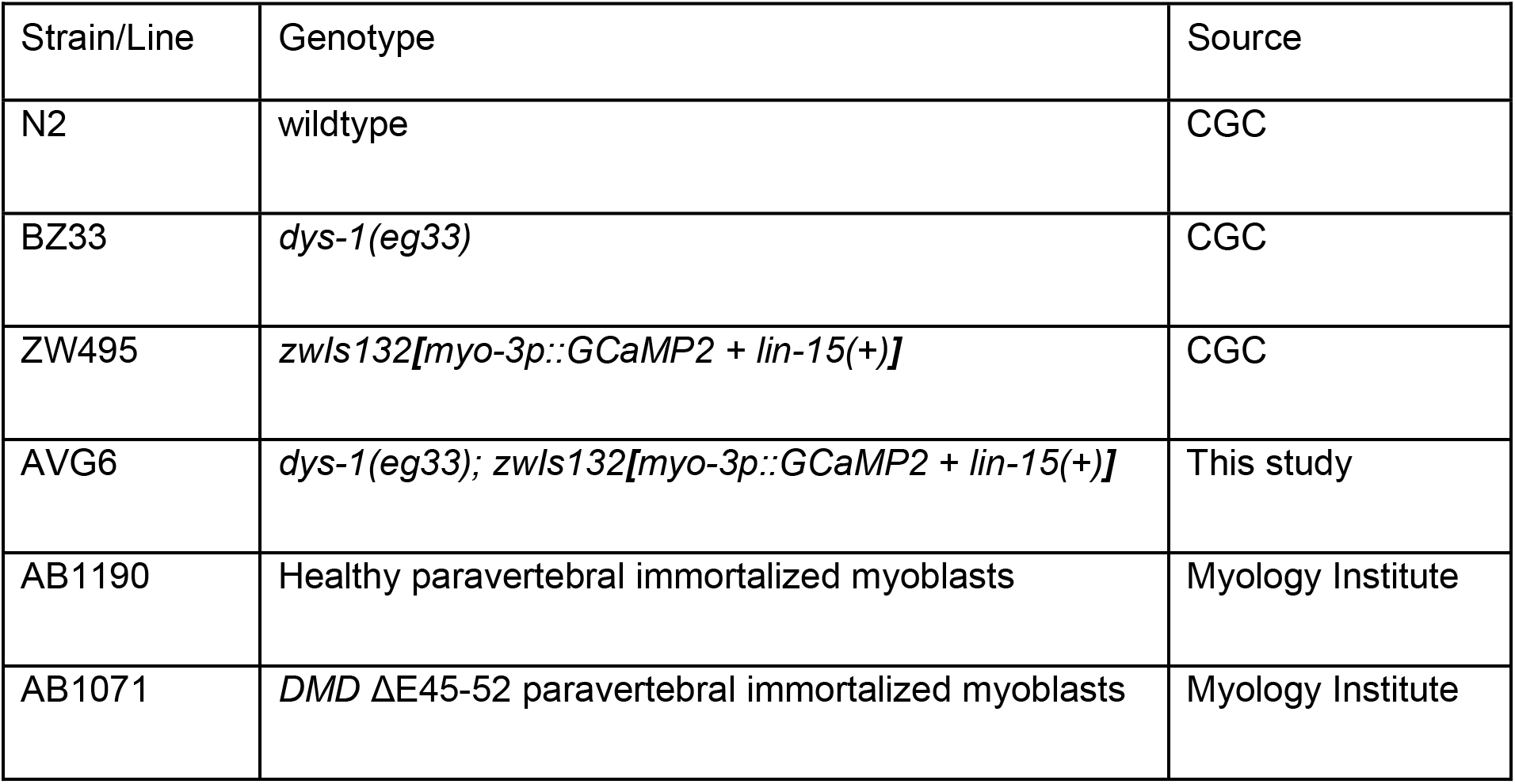
*C elegans* strains and human cell lines used in this work.

| Strain/Line | Genotype | Source |
|---|---|---|
| N2 | wildtype | CGC |
| BZ33 | *dys-1(eg33)* | CGC |
| ZW495 | *zwIs132**[**myo-3p::GCaMP2 + lin-15(+)**]*** | CGC |
| AVG6 | *dys-1(eg33); zwIs132**[**myo-3p::GCaMP2 + lin-15(+)**]*** | This study |
| AB1190 | Healthy paravertebral immortalized myoblasts | Myology Institute |
| AB1071 | *DMD* ΔE45-52 paravertebral immortalized myoblasts | Myology Institute |

